# Methodological choices matter: A systematic comparison of TMS-EEG studies targeting the primary motor cortex

**DOI:** 10.1101/2024.03.08.583873

**Authors:** Mikkel Malling Beck, Marieke Heyl, Louise Mejer, Mikkel C. Vinding, Lasse Christiansen, Leo Tomasevic, Hartwig Roman Siebner

**Affiliations:** Danish Research Centre for Magnetic Resonance, Centre for Functional and Diagnostic Imaging and Research, Copenhagen University Hospital - Amager and Hvidovre, Copenhagen, Denmark, Kettegård Allé 30, 2650 Hvidovre, Denmark; Department of Neuroscience, Faculty of Health and Medical Sciences, University of Copenhagen, Blegdamsvej 3B, 2200 Copenhagen N, Denmark; Department of Neurology, Copenhagen University Hospital Bispebjerg and Frederiksberg, Bispebjerg Bakke 23, 2400 København NV, Denmark; Department of Clinical Medicine, Faculty of Health and Medical Sciences, University of Copenhagen, Blegdamsvej 3B, 2200 Copenhagen N, Denmark

## Abstract

**Background:** Transcranial magnetic stimulation (TMS) triggers time-locked cortical activity that can be recorded with electroencephalography (EEG). Transcranial evoked potentials (TEPs) are widely used to probe brain responses to TMS.

**Methods:** Here, we systematically reviewed 137 published experiments that studied TEPs elicited from TMS to the human primary motor cortex (M1) in healthy individuals to investigate the impact of methodological choices. We scrutinized prevalent methodological choices and assessed how consistently they were reported in published papers. We extracted amplitudes and latencies from reported TEPs and compared total cortical activation and specific TEP peaks and components.

**Results:** Reporting of methodological details was overall sufficient, but some relevant information regarding the TMS settings and the recording and pre-processing of EEG data were missing in more than 25% of the included experiments. The published TEP latencies and amplitudes confirm the ’prototypical’ TEP waveform of M1, comprising distinct N15, P30, N45, P60, N100, and P180 peaks. However, variations in amplitude and latencies were evident across studies. Higher stimulation intensities were associated with overall larger TEP amplitudes. Active noise masking during TMS generally resulted in lower TEP amplitudes compared to no or passive masking but did not specifically impact those TEP peaks linked to long-latency sensory processing. Studies implementing independent component analysis (ICA) for artifact removal generally reported lower TEP amplitudes.

**Conclusion:** Some aspects of reporting practices could be improved in TEP studies to enable replication. Methodological choices, including TMS intensity and the use of noise masking or ICA, introduce systematic differences in reported TEP amplitudes. Further investigation into the significance of these and other methodological factors and their interactions is warranted.

## 1. Introduction

Transcranial magnetic stimulation (TMS) induces synchronized activity in cortical principal cells by directly stimulating neurons, leading to the generation of action potentials and the propagation of neural activity through connected networks [1]. Electroencephalographic (EEG) recordings can capture the synchronized activity evoked by TMS in cortical principal cells, because the synchronized polarization of these cells caused by postsynaptic potentials contribute to the electrical fields that can be detected on the scalp [2]. The synchronous activity generated by TMS can be traced in the averaged EEG as transcranial evoked potentials (TEPs) with characteristic positive and negative deflections (i.e., peaks) time-locked to the time of stimulation [3]. The TEP is a compound signal reflecting a mixture of local and remote cortical responses to TMS that provides valuable insights into the reactivity and connectivity of the human cortex at high temporal resolution [4].

Since the first studies demonstrated the possibility of concurrent TMS and EEG [5,6], the number of studies combining TMS with EEG recordings has increased steadily. The upsurge in popularity and the increased availability of the method can pave the road for important new insights into the underlying physiology of the TEP, and thereby how TMS ‘engages’ the brain. This may also add to the utility of TMS-EEG as a tool not only in research, but also in the clinic [7]. However, it also warrants the need for clear best practice guidelines and standards in terms of data acquisition, preprocessing, and analysis. This is critical because the choice of specific data acquisition, preprocessing, and analysis methods may ultimately affect the TEPs that are reported and, therefore, also the physiological insights that can be drawn. For example, recent studies have suggested that the later TEP peaks (>60ms) following TMS likely reflect the secondary processing of sensory co-activation from the stimulation (e.g. from auditory and somatosensory inputs) rather than the direct cortical activation and that this effect can be minimized, e.g., by the use of noise masking [8–10]. It has also been demonstrated that adopting different preprocessing strategies of the acquired TMS-EEG data may lead to significantly different TEPs [11,12]. These examples showcase how the abundant methodological degrees of freedom that a researcher is faced with when planning, performing, and analyzing a TMS-EEG experiment can affect the outcome. However, a comprehensive overview of the use and impact of various methodological choices on TEPs is currently lacking. A prerequisite for understanding the impact of methodological choices is that they are sufficiently reported. However, the vast parameter space associated with TMS-EEG studies may lead to an unfortunate omission of certain methodological details.

Many TEP studies have investigated the cortical response profile of the primary motor cortex (M1), mainly by targeting the hand representation. The interest in M1 can be explained by the fact that it is easily accessible to TMS due to its relatively superficial location, and because TMS of M1 elicits motor evoked potentials (MEPs) in contralateral limb muscles when delivered at intensities above resting motor threshold. For TEPs, a single TMS pulse to the M1 gives rise to a response lasting up to approx. 300ms with a characteristic sequence of negative and positive deflections, including the N15, P30, N45, P60, N100, P180 and N280 peaks [13–16].

The aim of this systematic review is to provide a systematic and comprehensive overview of the methods used in published TMS-EEG papers targeting the M1 and how this affects the TEP. We start by examining how rigorously methods are reported in previously published papers. Then, we illustrate the variability of methodological choices between published studies in the field and how used methods have changed over time. Finally, we outline how chosen methods affect various features of the reported TEPs. We focus on TMS-EEG studies conducted in healthy humans and restrict the systematic review to studies with original data originating from stimulation of the M1.

## 2. Methods

### 2.1 Search strategy

Published studies were identified using searches in the PubMed and the Web of Science databases. Searches were performed on January 10^th^, 2023. Combinations of the following search terms were used: ‘TMS-EEG’, ‘transcranial magnetic stimulation’, ‘TMS’, ‘electroencephalography’, ‘EEG’, ‘motor cortex’, ‘primary motor cortex’, ‘M1’. Articles were retrieved and duplicates were removed using Mendeley (v1.19.8) software algorithms and visual inspection.

### 2.2 Article screening and selection

The screening of the abstracts and the full texts was performed independently by three of the authors (MMB, MH & LM). Studies were included if they were (1) peer-reviewed and in English, (2) included adult participants with no diagnosed neurological or psychiatric disorders, and (3) reported an original variant of a time-domain response to single-pulse TMS over M1 (i.e., TEPs, local mean field power (LMFP), global mean field power (GMFP), butterfly plot or similar). After reviewing the articles, a consensus meeting was held where results were compared, and it was decided whether articles could be included. Additional publications not discovered in the initial search were identified in the retrieved articles upon full-text review and added to the list of eligible articles if they fulfilled the inclusion criteria.

### 2.3 Data extraction

Various methodological variables were extracted from the included full texts. This included variables relating to the study sample (e.g., number of participants and their age), TMS stimulation (e.g., number of pulses, type of pulse waveform and stimulation intensity), experimental setup (e.g., use of noise masking or sham stimulation), EEG data acquisition (e.g., amplifier type as well as number and type of electrodes), EEG preprocessing (e.g., filters and ICA) and data reporting (e.g., time windows of figures, electrodes reported). If information on methodological variables was not explicitly reported in the full text or supplementary material, it was labeled as ‘Not reported’. For a full list of extracted variables, please refer to supplementary file S1. Please note that some methodological variables were subsequently further subdivided to reduce the number of levels. For example, for TMS intensity, we created three categories relative to the resting motor threshold from the various intensities used (subthreshold, threshold, and suprathreshold stimulation).

To outline the potential consequences of common methodological choices across our full dataset of included articles, we extracted latencies and amplitudes of TEPs, LMFPs, and GMFPs from figures in the papers using the web-application WebPlotDigitizer [17]. Specifically, figures were cropped and imported into WebPlotDigitizer. From here, x and y-axes were calibrated, and values were extracted by adding data points either at clear peaks within commonly specified time-windows of interest (N15: 10-20ms, P30: 20-40ms, N45: 40-55ms, P60: 50-70ms, N100: 70-150ms, P180: 150-240ms, N280: 240-350ms) or as close to center value of this time-window as possible, e.g. at 30ms for the P30 peak. The data extraction and coding of studies were performed independently by three of the authors (MMB, MH, and LM).

### 2.4. Data visualization and analysis

To provide an overview of the consistency of reported methods, we visualized the percentage of studies reporting (or not reporting) extracted methodological variables. To account for the fact that some methods are nested in another (e.g., information about specific algorithms and number of rounds of independent component analyses (ICA) is contingent upon ICA being performed), we added the number of studies eligible for each variable. We also visualized common methodological choices by plotting percentages of studies using specific methods in pie-donut charts. Please note that the figures contain information available when removing studies with missing information for the specific variable. For comparing the effects of methodological choices on TEP waveforms, we focused on a subset of methodological variables to reduce dimensionality, including stimulation intensity, use of noise masking, and use of ICA. These variables were chosen, because they represent key decisions that researchers make when planning, performing, and analyzing TMS-EEG experiments. Furthermore, they represented variables that were quite commonly reported across studies. To obtain a measure of total activation following TMS we computed a composite ‘activation Index’ by rectifying and summating the extracted TEP peak amplitudes. To visually display how used methods affected specific TEP peaks, the extracted latencies and amplitudes for each peak were plotted by category (e.g., stimulation intensity: subthreshold, threshold, or suprathreshold stimulation) and a locally estimated scatterplot smoothing (LOESS) regression line was fitted between the observations. We statistically compared the amplitudes between categories using one-way ANOVAs and unpaired t-tests or the equivalent non-parametric alternatives when model assumptions were not met. P-values from statistical tests were Bonferroni-corrected. The procedure outlined above was only performed for studies reporting TEPs where peak latencies and amplitudes could be extracted (i.e., studies reporting LMFPs or GMFPs were not included in this analysis). Finally, we used parallel sets diagrams to visualize potential dependencies between categorical methodological choices. Data visualization and statistical analyses were carried out in R (v. 4.1.2; 2021-11-01).

## 3. Results

### 3.1 Database search and article selection

A total of 1729 papers were screened for duplicates and by title and abstract. 1484 of these studies were excluded, which left 245 papers for full-text screening. 121 of these were excluded as they did not fulfil the inclusion criteria. The remaining 124 studies (reporting 137 distinct experiments in 1892 subjects) were included in the systematic review and entered in the qualitative and quantitative summaries. The selection process is summarized in a flow chart in Fig.1.

**Figure 1.**
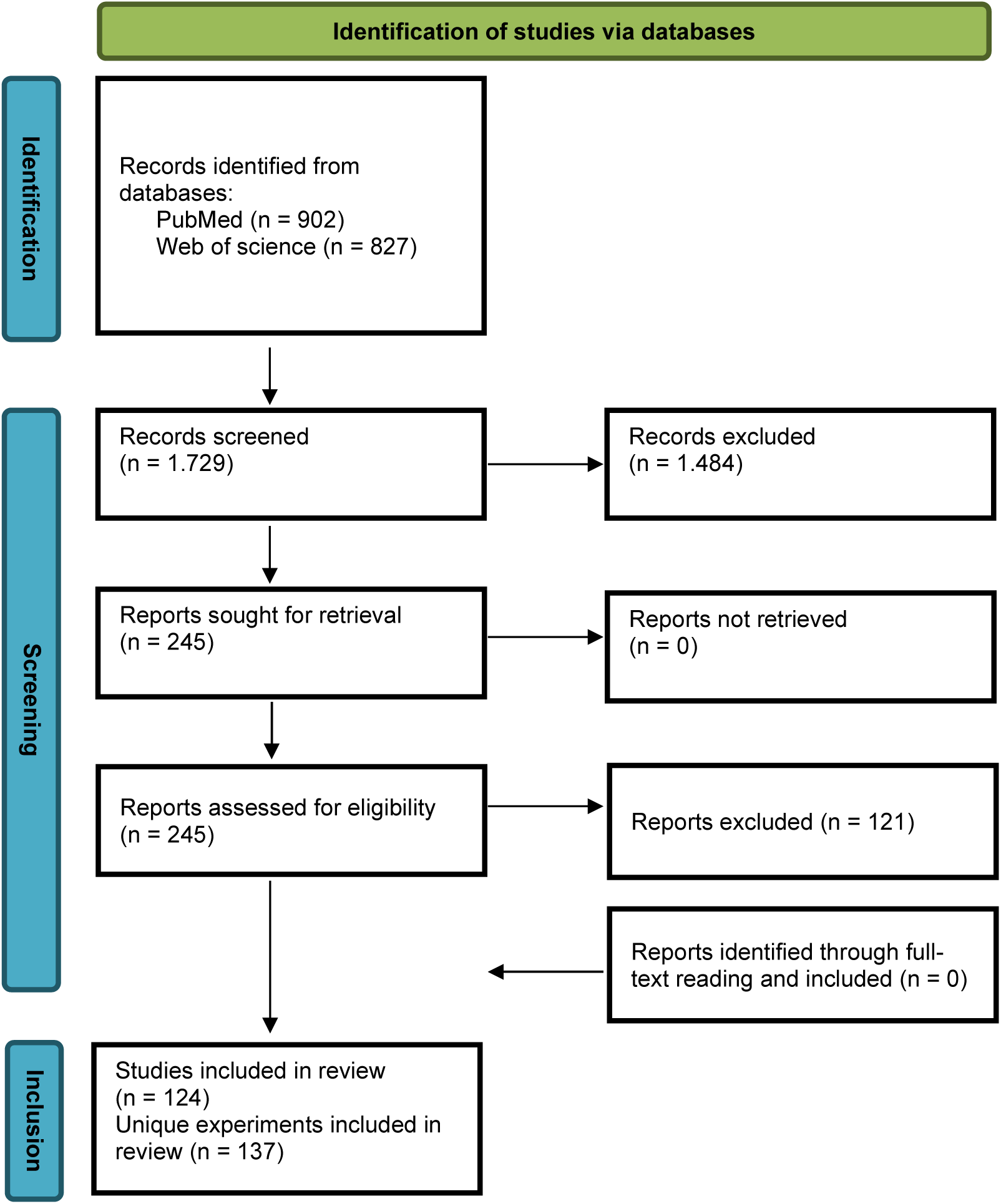
Flowchart of study inclusion.

### 3.2 Methodological reporting in published TMS-EEG studies

The frequency of reporting methodological variables varied substantially across published TMS-EEG studies. Figure 2 summarizes the reporting frequencies of methodological variables across six categories: the study demographics, TMS methodology, experimental setup, EEG acquisition, EEG preprocessing, and data reporting.

**Figure 2.**
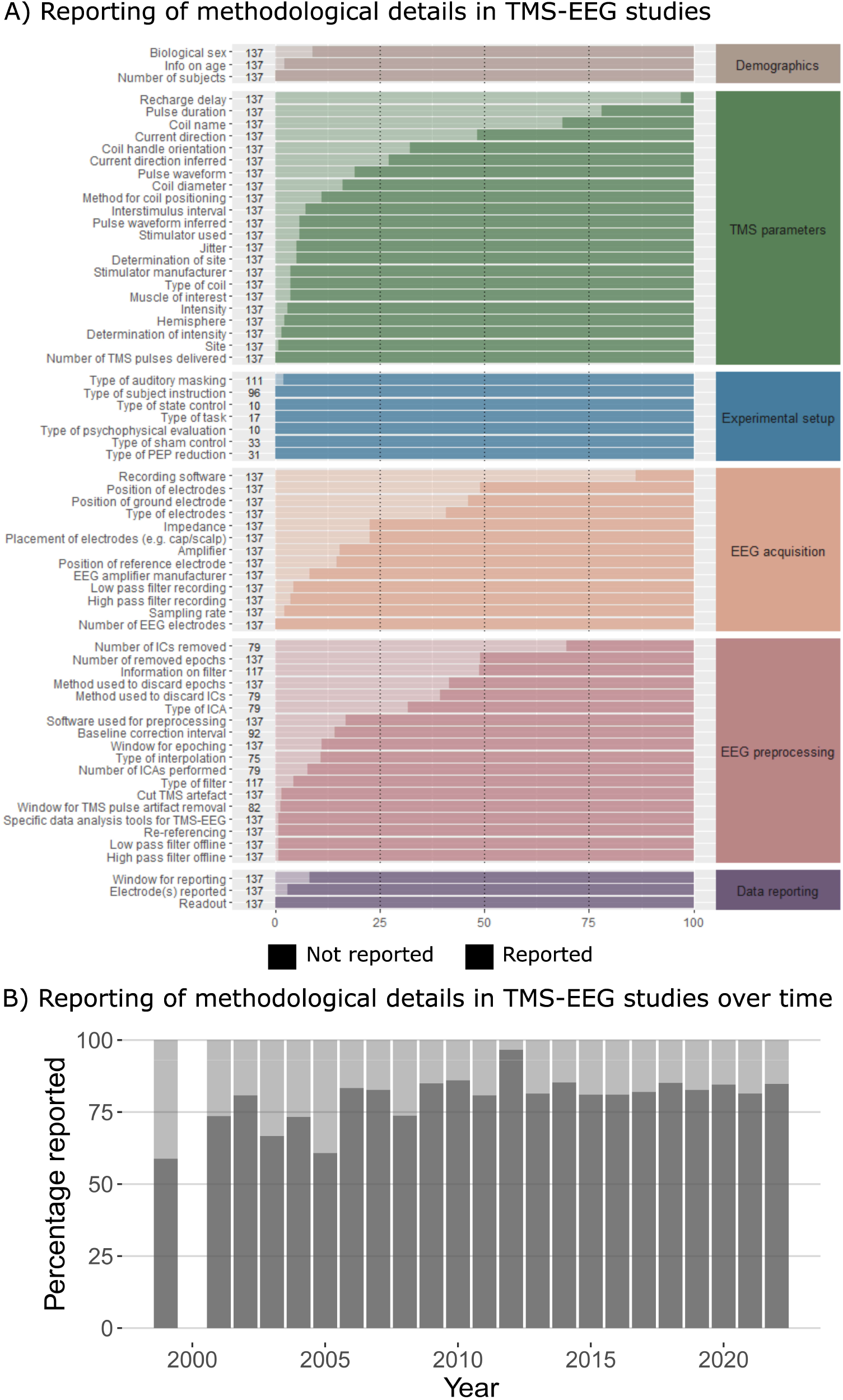
Reporting of methods in published TMS-EEG studies. (A) Percentage of reported (dark colors) and not reported (bright colors) information (x-axis) for various methodological variables (y-axis) divided into six methodological categories. (B) Percentage of reported methodological details (y-axis) across all variables over time (x-axis). After an initial increase, the percentage of reported variables has plateaued around 80% since 2006 with only minor fluctuations.

Information regarding the sample characteristics (e.g., number of participants, their biological sex, age), experimental set-up (e.g., auditory masking, experimental state, sham control), and data reporting (e.g., time window, EEG electrodes, EEG readout) were provided in more than 90% of the included studies (Fig. 2).

For TMS methodology specifically, more than 90% of the studies described the number of TMS pulses per TEP condition, the timing of consecutive TMS stimuli, the stimulation site, and how TMS intensity was determined. However, other relevant information regarding TMS methodology was reported in less than 75% of the included studies, such as the induced current direction, pulse duration, and recharge delay (see Fig. 2). For example, the direction of the electrical current induced in the coil or in the brain was only explicitly mentioned in ∼50% of studies and was only possible to infer from provided information in 72% of published studies.

Details relating to the EEG data acquisition, including listing the number of electrodes, sampling rate, band-pass filter settings, and the manufacturer of the EEG device were reported in more than 90% of the included studies. On the other hand, the type and position of recording electrodes, and the recording software were reported in less than 75% of the studies.

Level of reporting also varied for the procedures relating to preprocessing of the EEG data. For instance, the high-pass and low-pass cut-offs used for offline filtering, re-referencing, and the window for TMS-pulse artifact removal were specified in more than 90% of the publications. On the other hand, only 51% of the included studies reported the number of removed epochs and only 58% of the published studies clarified why these epochs were removed. Only 30% of TEP studies using ICA reported how many independent components were removed and only 60% disclosed the criteria on which ICs were discarded.

### 3.3 Common methodological choices and trends over time

Published TMS-EEG have used a wealth of different methods. In relation to the study sample, a median of 12 individuals participated in included TMS-EEG studies (Fig. 3A). Only eight studies had sample sizes above 25 with the number of participants in these studies ranging between 30 and 51. Most studies examined young adults. The median of the reported mean age was 28 (range: 21.0-77.8 years) and the median of the reported male-to-female ratio was approximately 3:2. Most studies were performed with subjects being at rest (88%) and only a few studies probed task-related differences in TEPs (12%), while participants performed motor (53%) or cognitive (47%) tasks (Figure S1).

**Figure 3.**
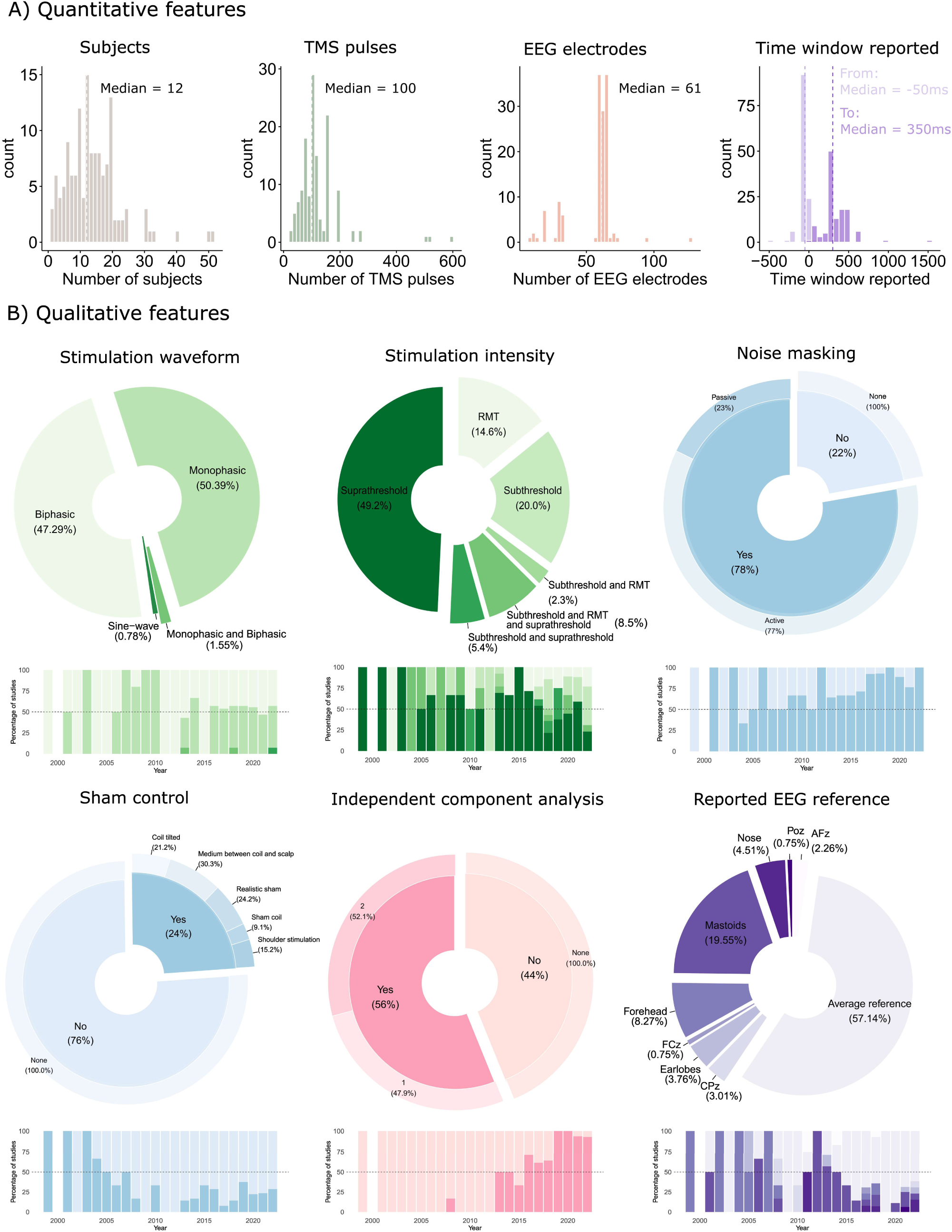
Common methodological choices and their development over time. The relative distribution of used methods in published TMS-EEG studies targeting the M1 and the percentage of studies using these methods over time are separately displayed for TMS parameters (green colors), acquisition parameters (blue colors), preprocessing steps (red colors), and data reporting (purple colors). (A) Each panel displays a specific ‘quantitative’ feature, including the distribution of number of included subjects, number of TMS pulses delivered, number of EEG electrodes used, and time window reported. (B) The pie-donut charts display the use of specific stimulation waveforms, stimulation intensities, noise masking, sham stimulation, independent component analysis, and the reported EEG reference. The corresponding bar-plots show the percentage of studies reporting the methodological variable per year, illustrating how methodological choices have developed over time.

Methodological choices regarding TMS varied considerably across studies. The median number of stimulations per TEP condition was 100, but many studies used much lower or higher number of TMS trials per TEP condition (Fig. 3A). The median interstimulus interval (ISI) between two consecutive TMS pulses was 4.75 sec (range: 1-12 sec) and 80% of the studies used jittered ISIs. TMS-EEG studies have almost equally often used monophasic and biphasic pulse waveforms (Fig. S1). Regardless of the waveform, 96% of studies that reports the induced current direction in M1 had the most influential phase of the TMS pulse induced in a posterior-to-anterior (PA) direction. Most TMS-EEG studies adjusted stimulation intensity based on individual resting motor threshold (85%), but TMS intensities varied across studies. 84% of the included papers only used a single TMS intensity, while the remaining 14% used multiple TMS intensities to investigate stimulus-response relationships. 74% of studies included a TMS intensity above resting motor threshold (RMT), while 36% of the studies included sub-motor threshold intensities and 26% of the studies included a TMS intensity that matched individual RMT (Fig. 2B). None of the methodological variables mentioned in this section seemed to show a systematic shift in its use over time.

78% of all included studies have used masking procedures to minimize the influence of auditory co-stimulation from the discharging coil (Fig. 3B), however masking procedures have differed across studies (Fig. 3B). ‘Active’ noise masking involves the administration of white or colored noise through earplugs or headphones and was more frequently used than ‘passive’ masking with earplugs or headphones (77% vs 23%). Direct coil-to-electrode contact was reported minimized in 23% of studies to reduce bone conduction of sound and vibratory somatosensory co-stimulation, by placing either a layer of foam (79%) or plastic (21%) between the coil and the electrodes (Fig. S1).

Sham TMS conditions that try to account for potential confounding effects from sensory processing have been used relatively rarely (24%). Most used sham protocols include introducing a medium between the coil and the head (such as air or wood), tilting the stimulation coil, stimulating the shoulder, and versions of a ‘realistic’ sham condition. While the two former sham protocols are primarily concerned with mimicking the auditory inputs from TMS, the two latter sham protocols are geared to mirror multisensory sensations associated with stimulation. However, perceived sensations of auditory and somatosensory experiences elicited by TMS were only reported in 7% of studies (Fig. S1).

EEG data were usually recorded using 64 scalp electrodes (Fig. 2A). Several different EEG amplifier systems have been used to acquire TMS-EEG data in TEP studies of M1. EEG recording systems include amplifiers with a large bandwidth that do not saturate from the electromagnetic TMS pulse (63%) or ‘sample-and-hold’ amplifiers that halt the acquisition around the delivery of the TMS pulse (37%), however this is rarely explicitly mentioned, and the proportion of large bandwidth amplifiers is likely greater based on the lower availability of ‘sample-and-hold’ amplifiers on the market. 59% of studies explicitly used an analogue high pass filter during data acquisition.

For preprocessing of TMS-EEG data, 75% and 82% of included articles performed offline high-pass and low-pass filtering, respectively. The most frequently used cut-off for the high-pass filter was 1Hz (53%) and for the low-pass filter it was 80Hz (56% of studies) (Fig. S1). 57% of included studies re-referenced their EEG data to the average of all electrodes (Fig. 3B). Other commonly used references were the mastoids (∼20%) and the forehead (∼10%). The TMS-artifact was cut out in 60% of studies and the modes of values used were -5 to 10ms (minimum: -20ms and maximum: 30ms). 55% of the studies used ICA to identify and deal with stimulation-related artefacts. The proportion of TMS-EEG studies using ICA has steadily increased over the last decade since its first use for artifact rejection in 2013 [18]. For the studies using ICA, 48% used a single round of ICA whereas 52% used two rounds of ICAs. The median of rejected independent components was 15, but this was rarely reported (30%) as mentioned above.

TEP data is most frequently reported from electrodes that are located closely to the site of stimulation, but data from the Cz electrode, i.e., the vertex, is also a relatively common choice (Fig. S2). The typically reported time window spans from -50 ms before to 350 ms after the administration of the TMS pulse (Fig. 3A).

### 3.4 Effects of methodological choices on TEPs

A curve fitted to the TEP latencies and amplitudes that were extracted from the included publications reproduced the ‘prototypical’ TEP waveform that has been reported in the literature for TMS of M1, consisting of distinct N15, P30, N45, P60, N100, and P180 peaks (Fig. 4A). The extracted data showed substantial variations of the individual TEP peaks across studies in terms of latency and amplitude (Fig. 4A).

**Figure 4.**
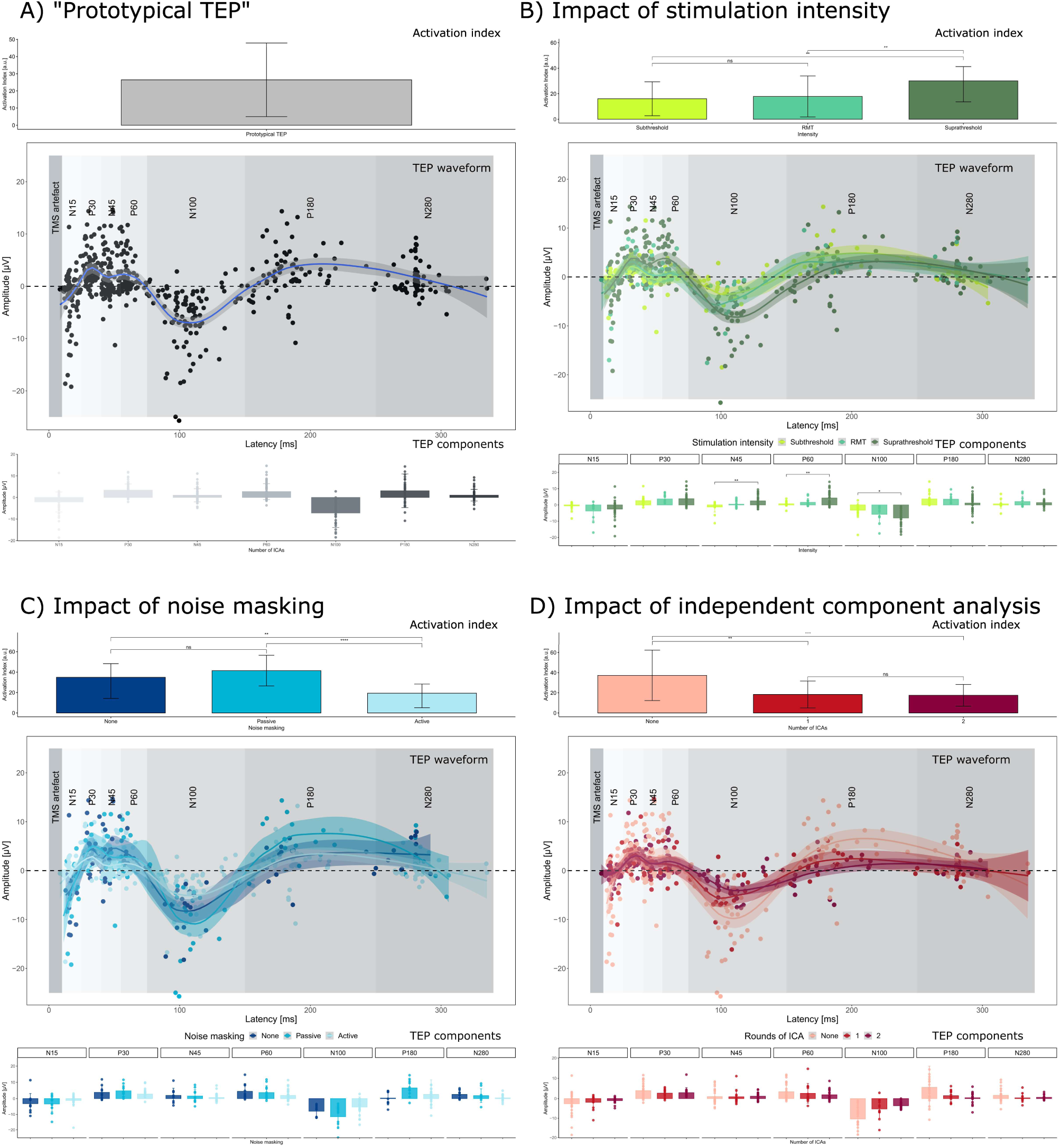
The impact of methodological choices on transcranial evoked EEG potentials. For each subfigure, the overall ’activation index’, computed as the sum of the absolute peak values for each TEP component, is presented in the top row. The middle row displays TEP waveforms reconstructed from extracted latencies (x-axis) and amplitudes (y-axis) and the bottom row displays a bar-graph of average values of different prototypical peak components for each of the methodological conditions. (A) Displays a ’prototypical TEP’ by combining all data irrespective of methodological choices. (B) Illustrates the impact of stimulation intensity all other parameters aside. Suprathreshold stimulation leads to a greater overall ’activation index’ across studies. The N45, P60 and N100 display greater amplitudes for suprathreshold stimulation compared to subthreshold and threshold intensities. (C) Displays the impact of noise masking all other parameters aside. Studies using active noise masking generally result in a smaller overall ’activation index’ and this effect seems to be general across various components. (D) Shows the impact of using independent component analysis for preprocessing all other parameters aside. Studies using ICA generally result in a smaller overall ’activation index’, and this seems to be general across multiple peak components, including peaks that are clearly physiological such as the N100.

We statistically evaluated the effects of methodological choices on reported TEPs by plotting and comparing extracted latencies and amplitudes of these prototypical peaks. We found a clear effect of TMS intensity on the reported TEPs (Fig 3B): TMS at an intensity above RMT displayed greater overall activation, as reflected by the TEP ‘activation index’, compared to subthreshold stimulation and stimulation at resting motor threshold (F_(2, 62)_=5.38, P=0.007). When comparing specific TEP peaks, the N45, P60, and N100 amplitudes were significantly larger in amplitude following suprathreshold stimulation compared to both subthreshold and threshold stimulation (Fig. 4B, bottom). Notably, no significant differences between TMS intensity were observed when comparing the N100-P180 complex (F_(2, 74)_=0.90, P=0.41). (Fig 5A). Together, the results indicate that higher stimulation intensities generally result in greater TEP amplitudes, especially of the N45, P60 and N100 peaks.

**Figure 5.**
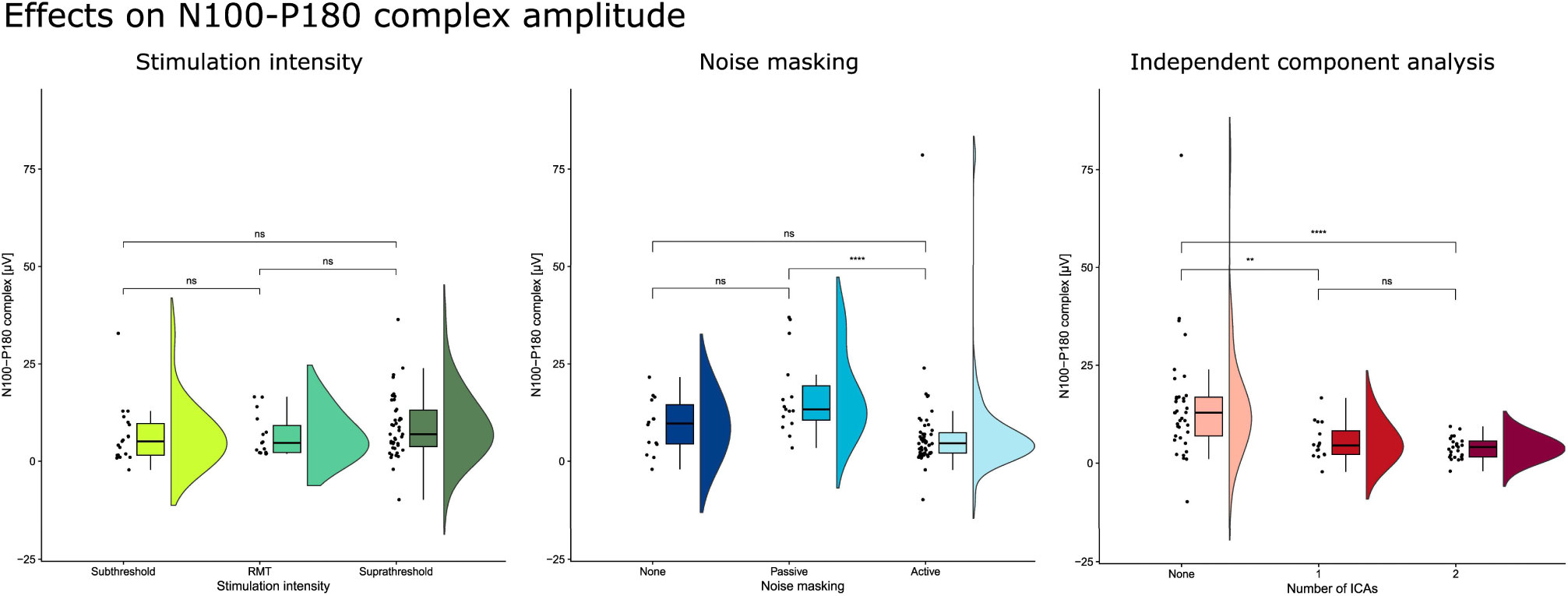
The impact of methodological choices on N100-P180 complex. Displays the effect on stimulation intensity (left), noise masking (middle), and independent component analysis (right) on reported N100-P180 peak-to-peak amplitudes.

We also compared the TEP data obtained in studies that used active or passive auditory masking to studies that did not apply auditory masking. A significant effect was observed on cortical activation, as reflected by the TEP ‘activation index’ (F_(2, 64)_=7.74; P<0.001). Active noise masking was associated with a lower activation index compared to both passive noise masking (e.g., earplugs) (t(53)=-3.35; P=0.002) and no noise masking (t(53)=-2.50; P=0.044, Fig. 4). Interestingly, when comparing specific TEP peaks, no significant differences were observed between studies after correction for multiple comparisons. When using the peak-to-peak amplitudes of the N100-P180 complex as a cortical marker of long-latency sensory processing, there was a significant main effect of the type of auditory masking (F_(2, 76)_=4.91, P=0.009, Fig 5B). Post-hoc testing revealed significant higher peak-to-peak amplitudes of the N100-P180 complex for studies using passive noise masking relative to active noise masking (t(64) =3.13; P=0.013). No effect was found between studies using passive and no noise masking (t(28)=2.39; P=0.074), or between active noise masking and no noise masking (t(63)=0.83; P=1). In sum, active noise masking was associated with a general reduction of TEP amplitudes but had no distinct effect on TEP peaks related to long-latency sensory processing.

Using 1 or 2 rounds of ICA during pre-processing of the EEG data resulted in a smaller overall ‘activation index’ compared to studies that did not use ICA (Fig 4D) (F_(2, 63)_=8.33, P<0.001). However, significant effects were only found for the N100 peak that was smaller when 2 rounds of ICA compared to not using ICA (t(44)=4.21; P=0.005). A significant effect was also observed when comparing the N100-P180 peak-to-peak amplitude (Fig 5C) (F_(2, 75)_=9.23, P<0.001). Collectively, this suggests that using ICA leads to lower TEP amplitudes across various peaks, and particularly for specific TEP indices of long-latency cortical processing of sensory input (N100, N100-P180).

For all variables listed above, no significant differences were found between compared conditions for spontaneous baseline activity, i.e., before the stimulation (Fig S3), suggesting that the use of different methods did not result in overall shifts in spontaneous (pre-stimulus) EEG activity. Furthermore, comparable results between studies were obtained when comparing TEP features weighted by the relative sample size (Fig S4).

## 4. Discussion

Our systematic review of 137 published TMS-EEG experiments targeting the M1 revealed three main findings. First, the reporting of methodological details varied across studies. Second, the published data in healthy individuals converged on a ’prototypical’ TEP waveform of M1, comprising distinct N15, P30, N45, P60, N100, and P180 peaks despite the vast methodological diversity and variations in amplitude and latencies across studies. Third, we found that methodological choices including the intensity of TMS, the type of noise masking, and the use of ICA for artifact rejection introduce systematic differences in reported TEP amplitudes.

### 4.1 Methodological reporting in published TMS-EEG studies

Rigorous reporting of methodological variables is a prerequisite for the interpretation and replication of studies. In this review, we found that reporting of methods was overall good and had improved over time. We also found that many methodological variables were consistently reported in published studies, whereas some variables were not. The impact of these omissions cannot be inferred from this review. Some may be more critical than others. For example, missing information on the number of independent components that were removed following ICA, and the criteria that were applied to label these components as artefactual, may complicate interpretations of the data because the removal of components can significantly impact the reported TEP waveforms as shown here and in a study specifically investigating this question [11]. There is a continued need to make decisions related to data acquisition, preprocessing, and analysis fully transparent. This is crucial for ensuring the reproducibility of TMS-EEG experiments and analyses.

### 4.2 Methodological diversity and robustness of the TEP

When fitting a curve to the TEP latencies and amplitudes that we had extracted from all the included publications irrespective of the methods used in these studies, we were able to reproduce what is considered the ‘prototypical’ TEP waveform after stimulating M1: The characteristic time-locked response consisted of negative peaks occurring at around 15, 45 and 100 ms and positive peaks at around 30, 60 and 180 ms after transcranial stimulation [13–15]. In the light of the various sets of methods that have been applied in published studies to date, we infer that the EEG pattern that can be evoked by TMS of M1 is relatively robust in the healthy adult brain. At the same time, there was also some degree of variability across studies when comparing the actual latencies and amplitudes. Grouping the included studies by used methods further revealed how methodological choices can systematically affect TEP peak amplitudes. In what follows below, we will discuss how three key decisions that researchers have to make when planning, performing, and analyzing TMS-EEG experiments can affect reported TEP peak amplitudes.

### 4.3 The impact of common methodological choices on TEPs

Studies using suprathreshold stimulation intensities generally evoked larger cortical responses, as reflected by the ‘activation index’, compared to the TEPs that have been reported in studies using stimulation intensities at or below resting motor threshold. Additionally, significant differences between stimulation intensities were seen for specific TEP peaks. The N45, P60 and N100 amplitudes were significantly greater following suprathreshold stimulation compared to subthreshold stimulation.

The larger TEP response following suprathreshold TMS pulses can be attributed to a stronger transcranial excitation of the cortical circuits in the targeted M1. However, other “non-transcranial” off-target mechanisms may also contribute. For example, TMS-EEG studies using suprathreshold stimulation also elicit twitches in the target muscle, for instance in contralateral intrinsic hand muscles in the case of TMS-EEG studies targeting the motor hand representation. The muscle twitch generates re-afferent activity which reaches the somatosensory cortex contralateral to the twitch (i.e., ipsilateral to the stimulation site) approximately ∼40-60 ms after stimulation. This reafferent cortical excitation may contribute to larger N45 and P60 peaks of the TEP in the case of suprathreshold stimulation as opposed to subthreshold stimulation. This hypothesis is supported by a study that stimulated at resting motor threshold and compared TEPs from trials where the transcranial stimulus elicited a motor response to TEPs from trials where the same TMS pulse failed to produce a motor response [19]. Greater amplitudes were observed around this time window for trials that elicited MEPs compared to trials that did not.

The TEPs is a compound signal reflecting a mixture of local and remote cortical responses to TMS caused by “direct” transcranial excitation and “indirect" peripheral excitation of the nervous system [3,8]. Peripheral co-stimulation cannot be avoided and results in a complex “off-target” activation of cortical areas that contributes and interacts with the cortical response patterns caused by transcranial cortical stimulation [1]. A prominent “peripheral” response is caused by co-stimulation of the auditory system through air and bone conduction [1]. TEP peaks that have latencies of more than 60 ms have often been ascribed to processes related to late processing of auditory and somatosensory co-stimulation [8,9]. Since higher stimulation intensities are accompanied by louder click sounds [20] and a greater electrical excitation of neural elements outside the brain, a stronger peripheral off-target stimulation may contribute to a larger N100 peak at suprathreshold intensities. Yet, no clear effects were observed on the N100-P180 complex.

For the reason outlined above, noise masking is used to minimize potential contamination of TEP responses by cortical processing of the sound generated by the discharging coil [3,15]. Auditory stimuli give rise to a clear time-locked EEG response referred to as auditory evoked potentials (AEPs)[21]. AEPs are multifaceted responses, but within the TMS-EEG field, the focus has mainly been on late AEP components including a positive peak around 50ms (P50) and the subsequent N100-P180 complex [22]. As seen in the prototypical TEP depicted in Figure 4A, similar peaks are also observed after TMS of M1. When exploring the effects of noise masking on TEPs from previously published studies, we found an overall lower evoked cortical activation, as reflected by the ‘activation index’, in those studies using active noise masking (e.g., white noise delivered through earplugs) compared to studies that had used passive (e.g., earplugs) and no noise masking. This finding is interesting in the light of recent studies showing that white noise may change cortical excitability [23,24].

The effect of noise masking appeared to impact the TEP broadly rather than selectively influencing specific TEP peaks. No single TEP peak was found to be uniquely influenced by the type of noise masking. However, lower amplitudes of the N100-P180 complex, a hallmark signature of late cortical sensory processing, were observed when comparing studies applying active or no noise masking to studies passive noise masking. Of note, no consistent differences were observed when comparing the TEP responses in studies using active noise masking versus no noise masking. One may be tempted to jump to the wrong conclusion that noise masking is not effective in reducing auditory co-stimulation. When applied in a rigorous manner, the efficacy of active noise masking is convincing [25]. We rather attribute the absence of a consistent difference to insufficient use of masking procedures. Indeed, 6/6 of the studies included in this work that have assessed perceived loudness of the TMS “click” sounds report some degree of audibility of the stimulation despite using noise masking. This could be suggestive of a more general trend that previous attempts to mask the sound of the discharging TMS coil—passive as well as active—may have often been inefficient and that residual auditory input from stimulation may have contributed to the reported TEP results in many published TMS-EEG studies [26].

In this context, we would like to stress that the N100-P180 complex is not exclusively related to auditory processing, but also reflects a neural response to salient, modality-independent sensory stimuli [8,9]. For example, a clear N100-P180 response was observed in a deaf individual, when stimulation was delivered in proximity to the scalp [25]. In this case, the N100-P180 was most likely produced by stimulation-induced excitation of skin receptors and peripheral axons contributing to somatosensation [27]. Importantly, no N100-P180 response was seen when the coil was positioned further away from the head, effectively eliminating the somatosensory stimulation by the TMS pulse [25]. A systematic evaluation of the influence of subjective percepts of auditory and somatosensory co-stimulation on TMS-evoked cortical activity is hampered by the fact that only 7% of the reviewed studies assessed how participants perceived TMS (either TMS *per se* or as compared to sham). Another approach to deal with sensory co-stimulation is to adopt sham procedures to disentangle direct responses to TMS from responses resulting from sensory co-activation [8–10]. Although sham procedures are increasingly included in the experimental design, it still remains challenging to effectively match the perceived sensations between real and sham TMS [8,28,29]. Regardless of the inclusion of a sham control condition, we suggest that future TEP studies should routinely assess and report the multimodal sensory percepts elicited by the TMS pulse.

ICA is a method for separating multivariate data into statistically independent components and was originally introduced to the TMS-EEG domain as a tool for separating neurophysiological data from artefacts, such as the TMS pulse decay and scalp muscle artefacts [18]. We found that studies employing ICA generally reported smaller TEP peak amplitudes compared to those not using ICA. This overall suppressive effect of ICA generalized across many TEP peaks, indicating that the use of ICA may introduce a general bias towards the reported TEP dynamics in the cleaned signal. The impact of ICA on reported TEP amplitudes was most pronounced for the N100 peak and the N100-P180 complex. This shows that the use of ICA can selectively - but likely unintentionally - modify the expression of distinct physiological cortical signals inherent in the TEP. Our findings raise concerns about unintended co-removal of both physiological and artifactual data [30]. Indeed, the lower amplitudes observed in late TEP peaks, such as the N100, may indicate an inadvertent impact on distinct physiological signals. We see a need for a critical evaluation of the use of ICA in artifact removal in TMS-EEG studies. This becomes particularly relevant due to the growing popularity of ICA as over 90% of the reviewed TMS studies incorporated ICA in the last five years. Furthermore, many of these studies did not provide a detailed account of *how* ICA was used (i.e., the number of components and their selection) which hampers the reproducibility.

### 4.4 Concluding remarks

In the present work, we provide a comprehensive review of the different methods used in TMS-EEG research targeting the M1. We also show how reporting standards varied across studies and that some relevant details regarding the TMS procedures and recording and pre-processing of EEG data should be reported more consistently in future studies. We outlined how certain methodological choices impact TEPs by comparing extracted TEP amplitudes from previously published studies using different methods. We found that methodological choices regarding stimulation intensity, auditory masking, and the use of ICA are associated with systematic variations in the published TEP data. While all three methodological variables were associated with systematic variations in the published TEP data, we found no clear dependencies among the three methodological choices (Supplementary Fig. 5). These findings imply that methodological choices matter and should be carefully considered when comparing results between studies and when planning future studies. In this review, we focused on three specific methodological variables, namely stimulation intensity, the application of noise masking, and the use of ICA. These variables were chosen as they reflect critical considerations for researchers in the planning, execution, and analysis of TMS-EEG experiments. It can be assumed that other methodological variables may also exert systematic influences on TEP outcomes. The impact of methodological choices and their possible interactions warrant a systematic examination in future studies [31].

**Table 1.**
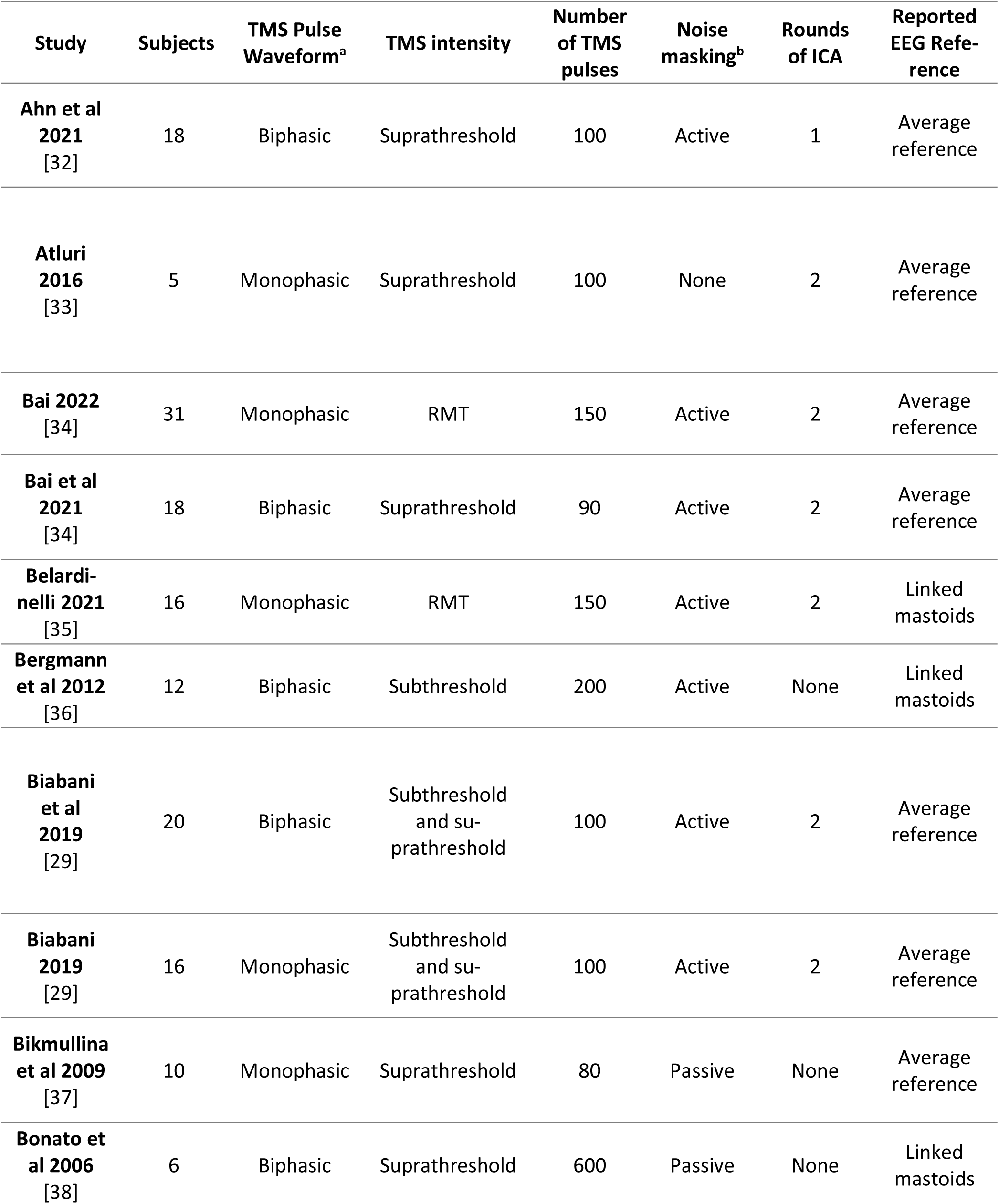
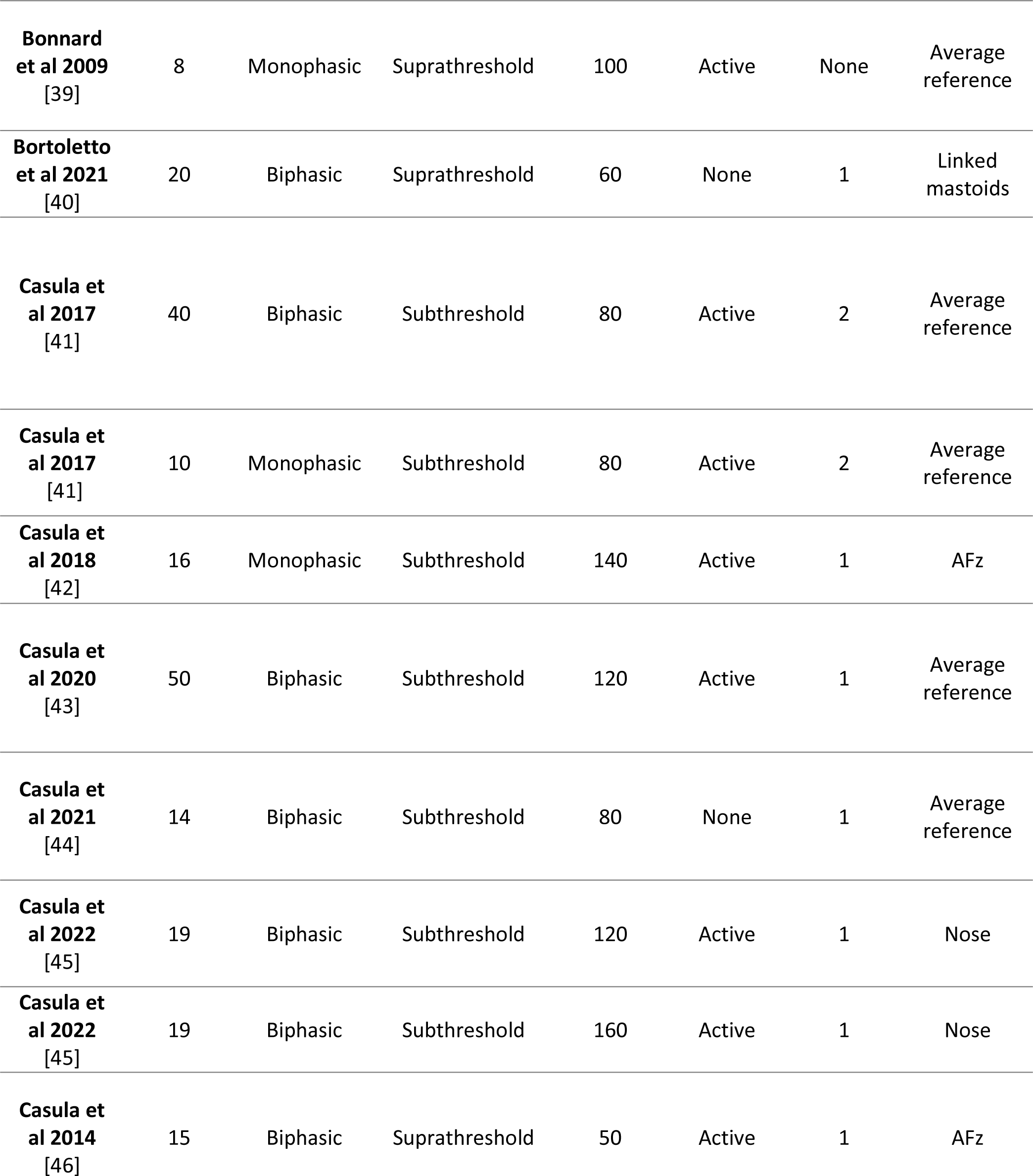
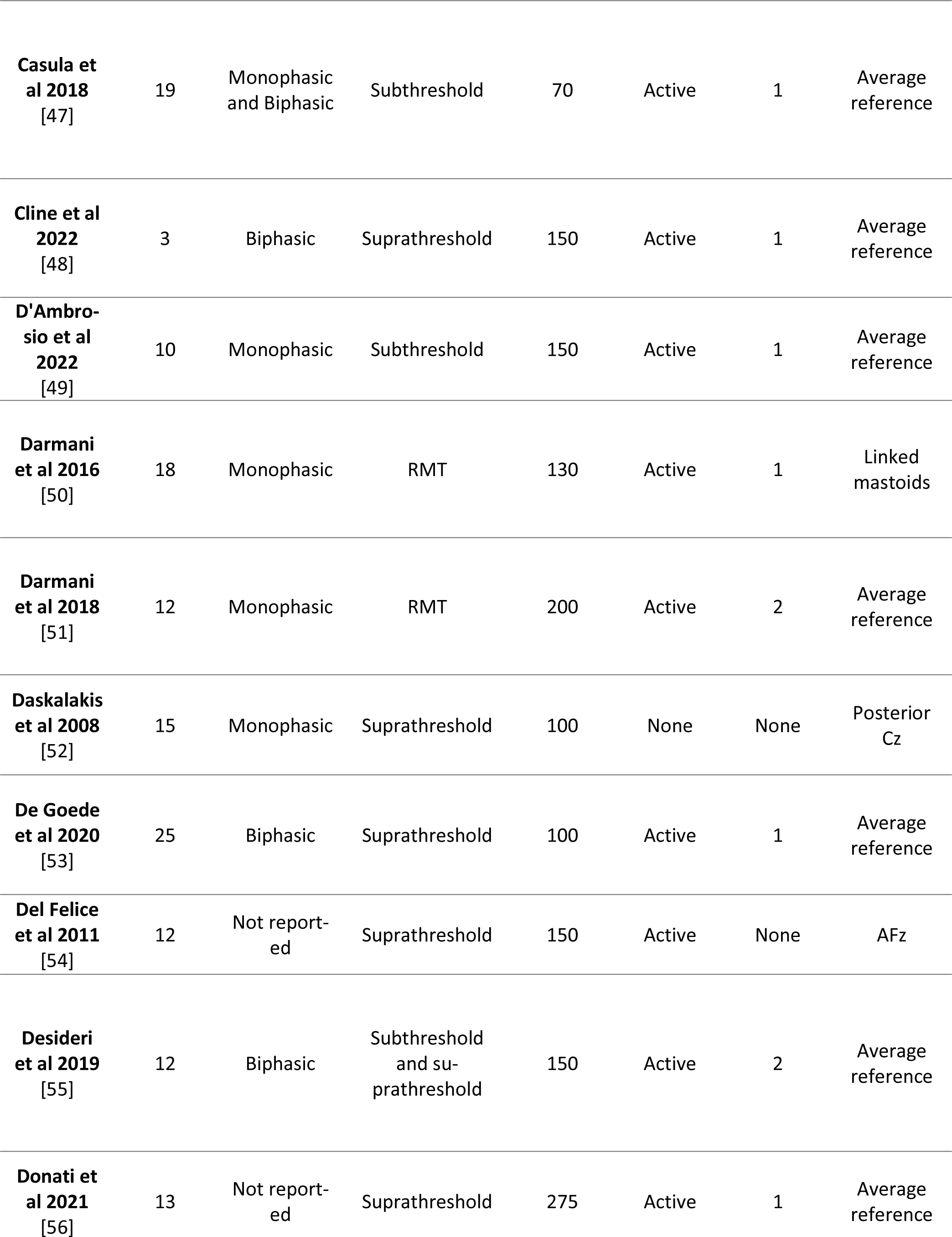
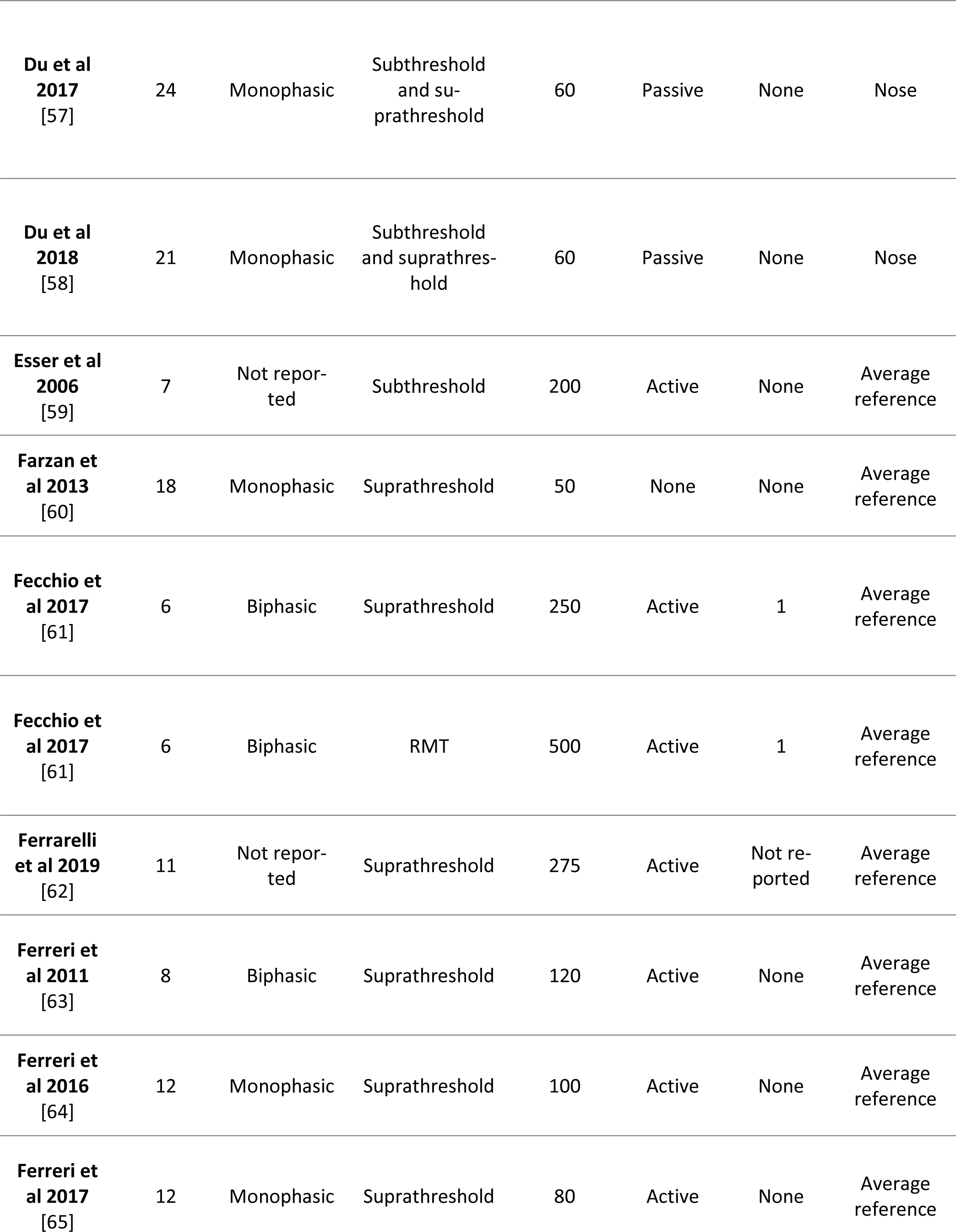
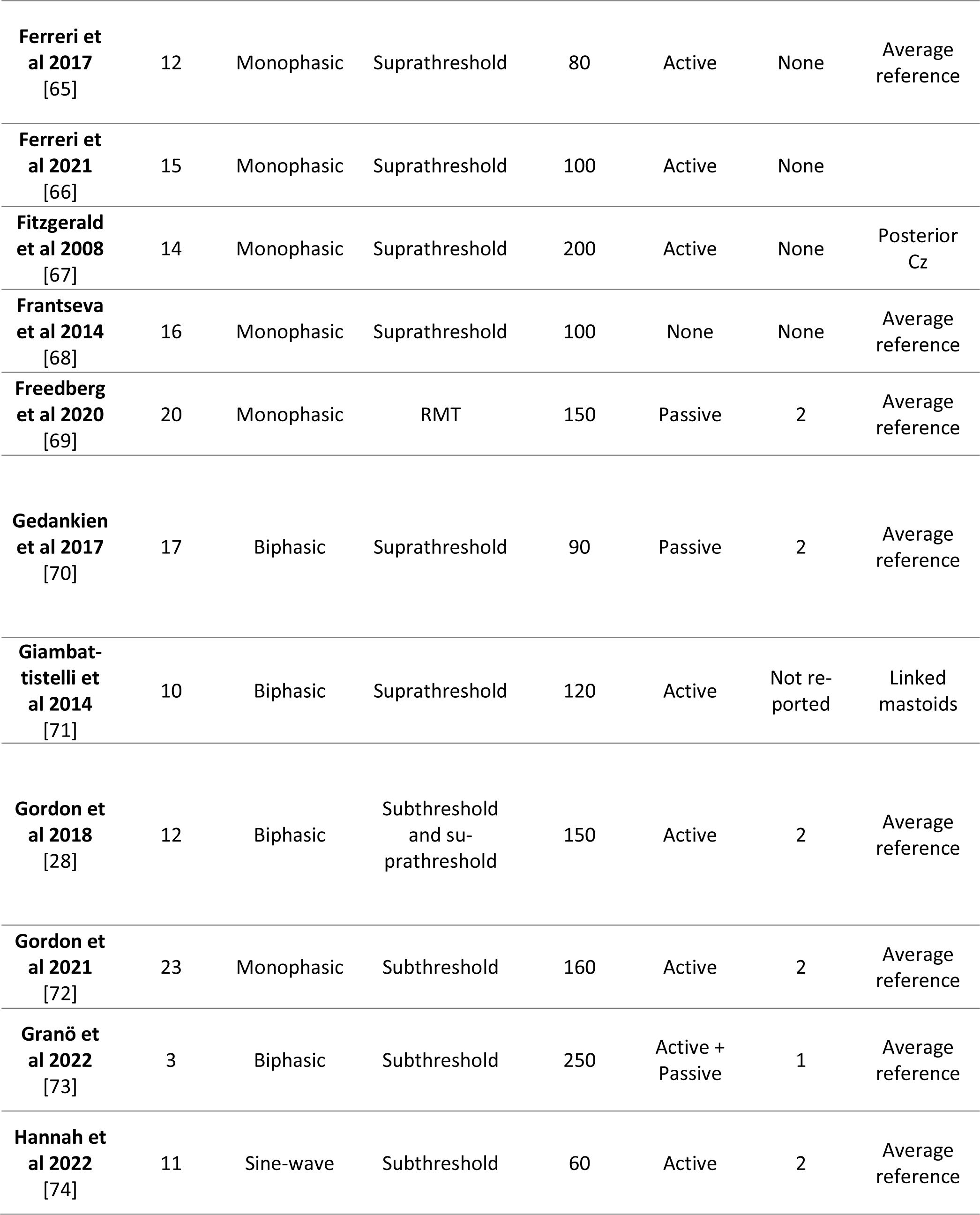
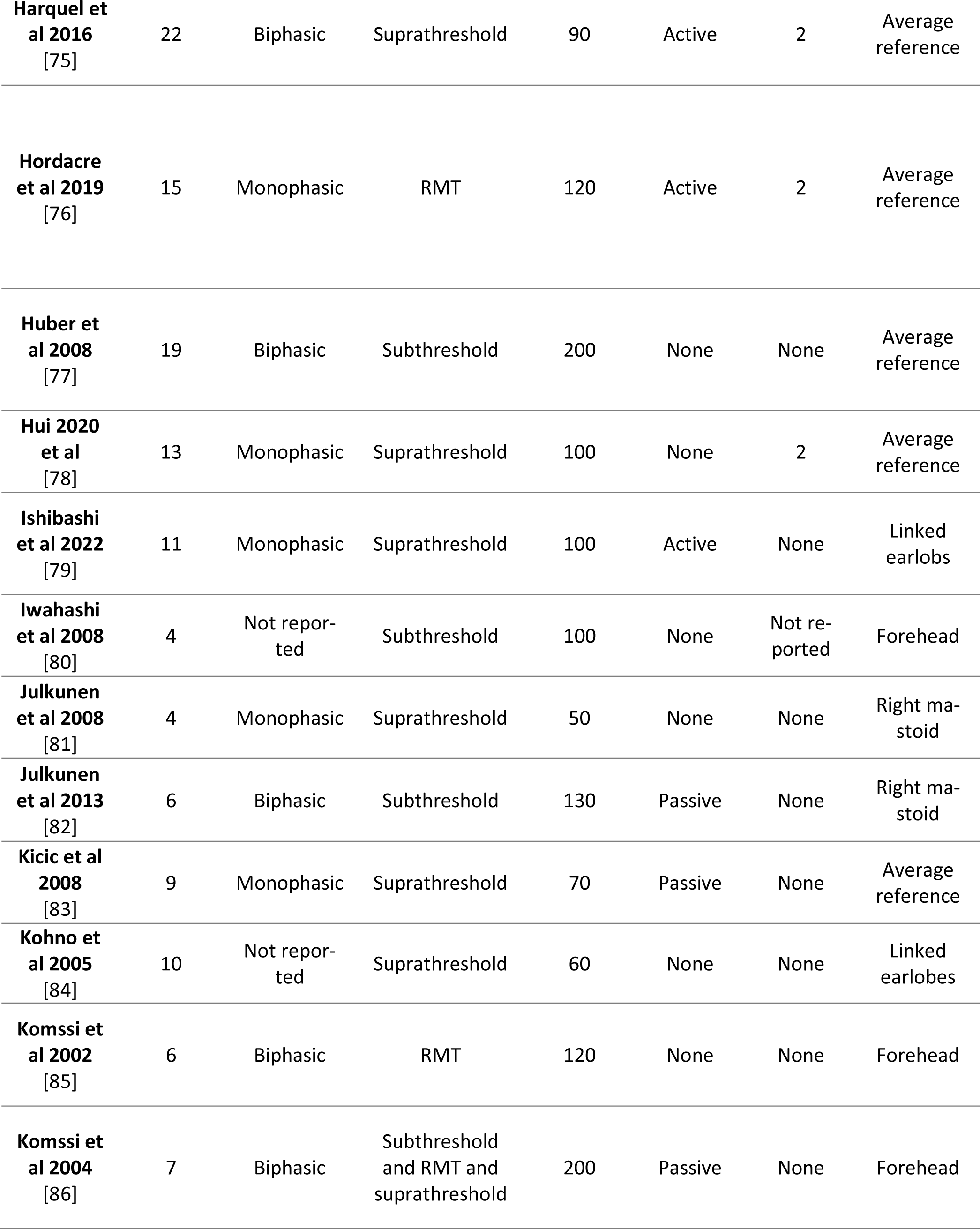
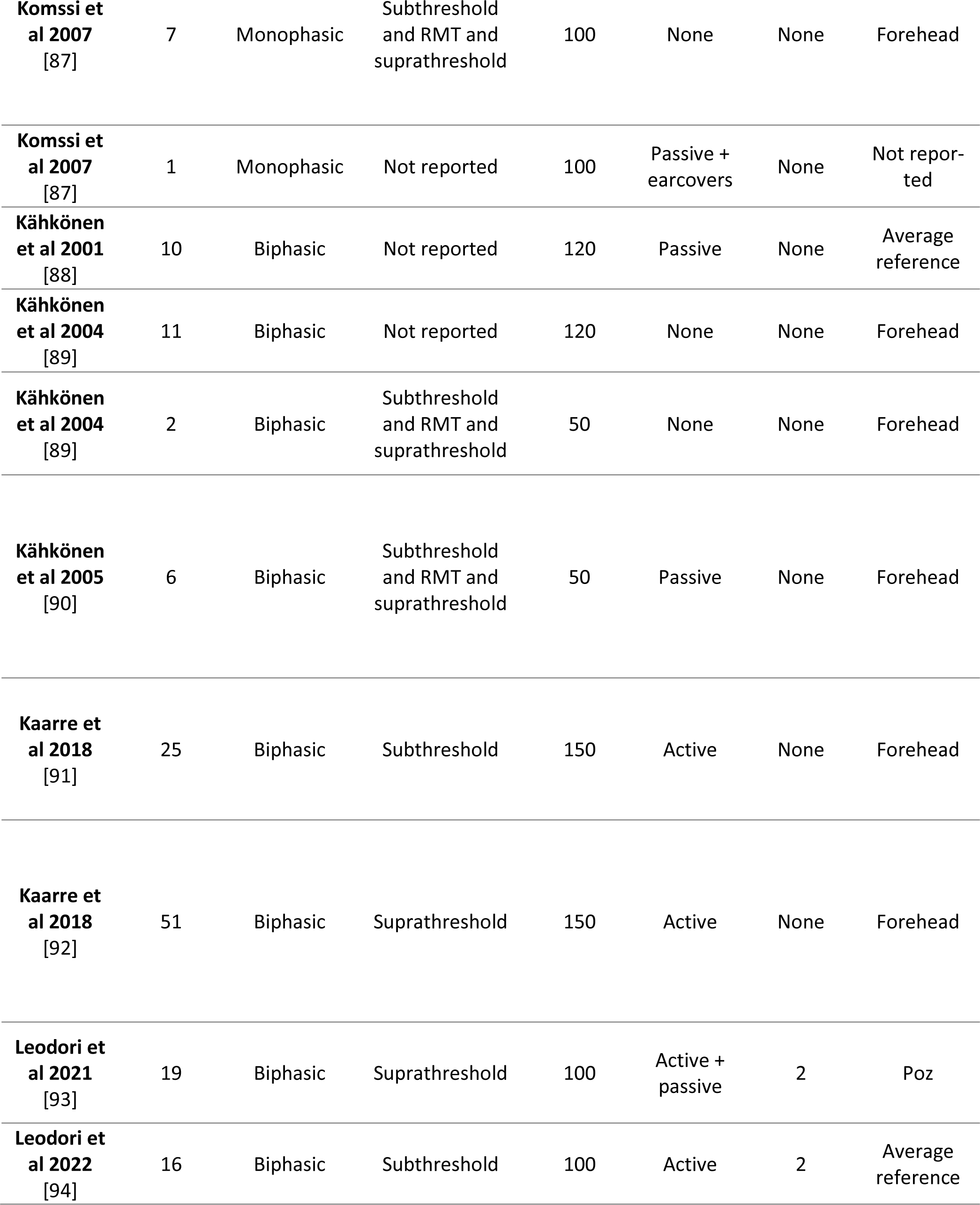
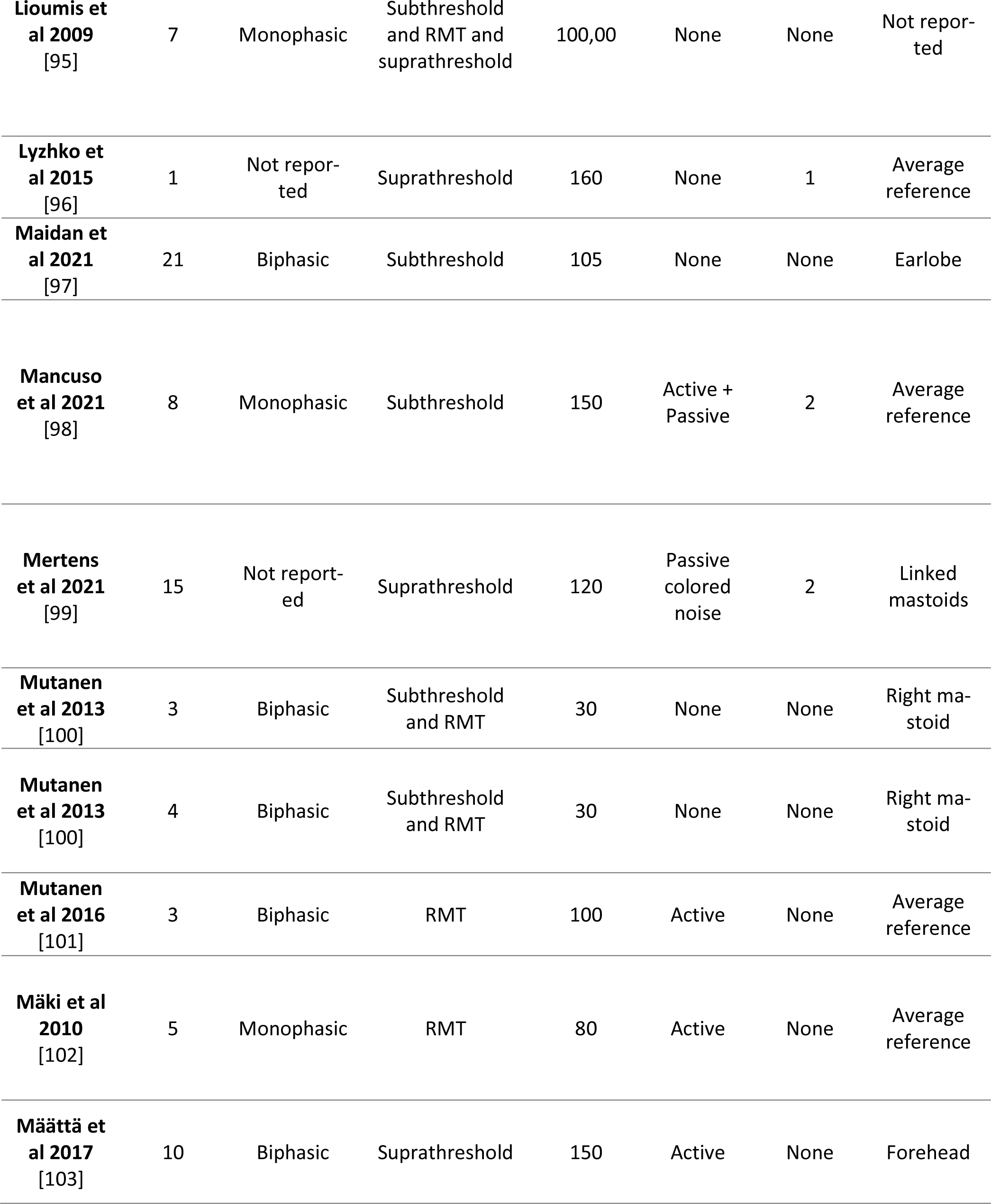
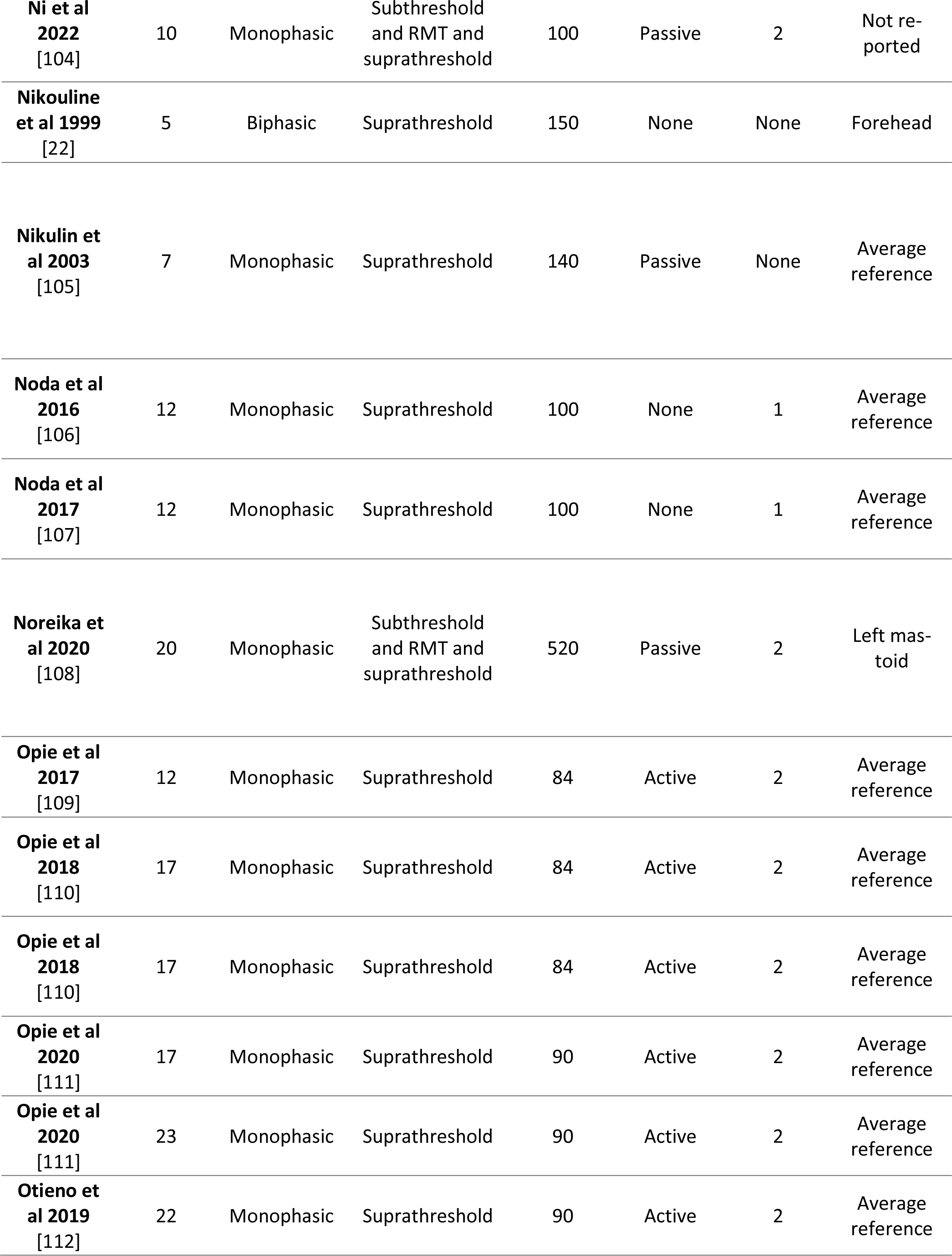
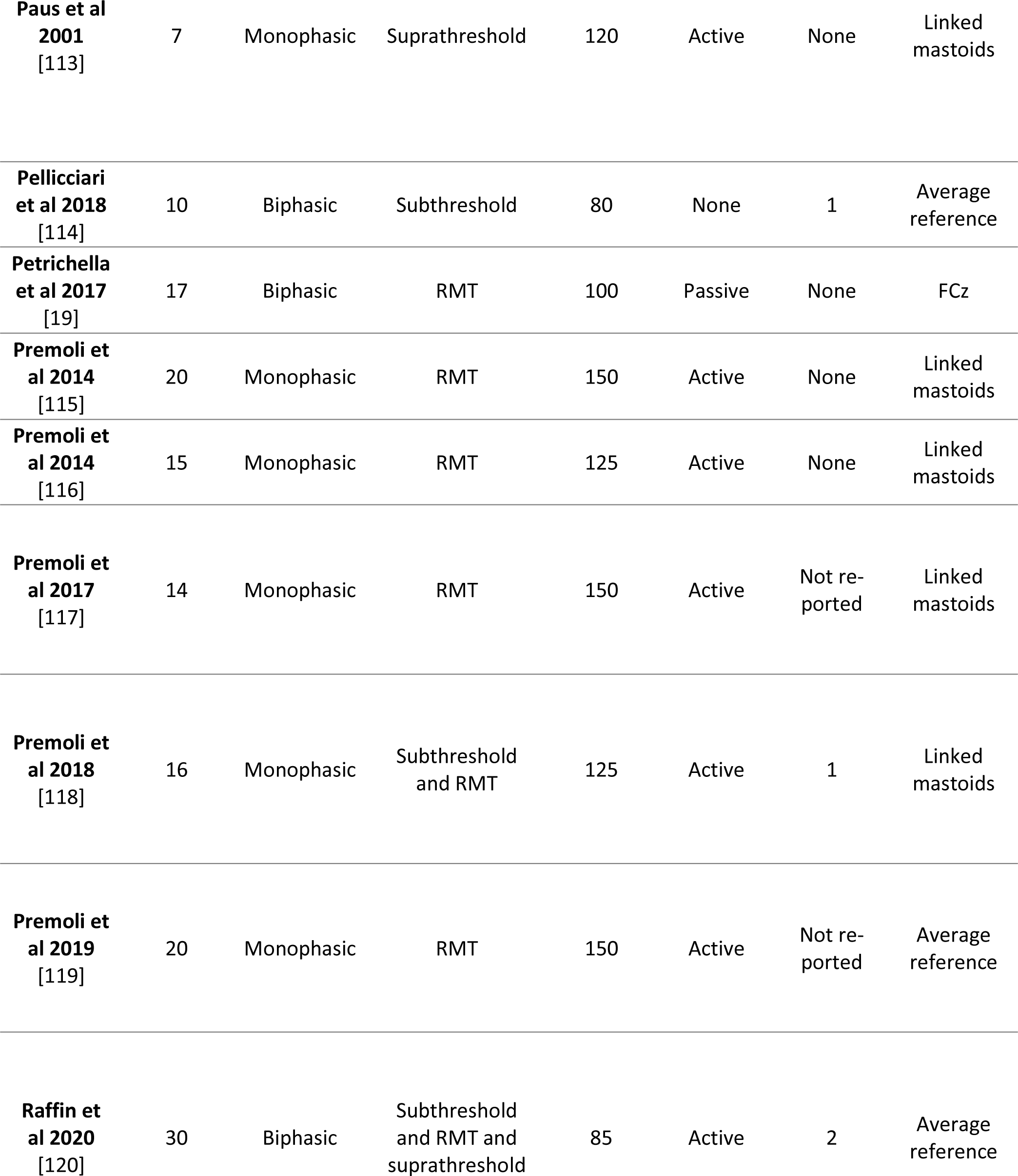
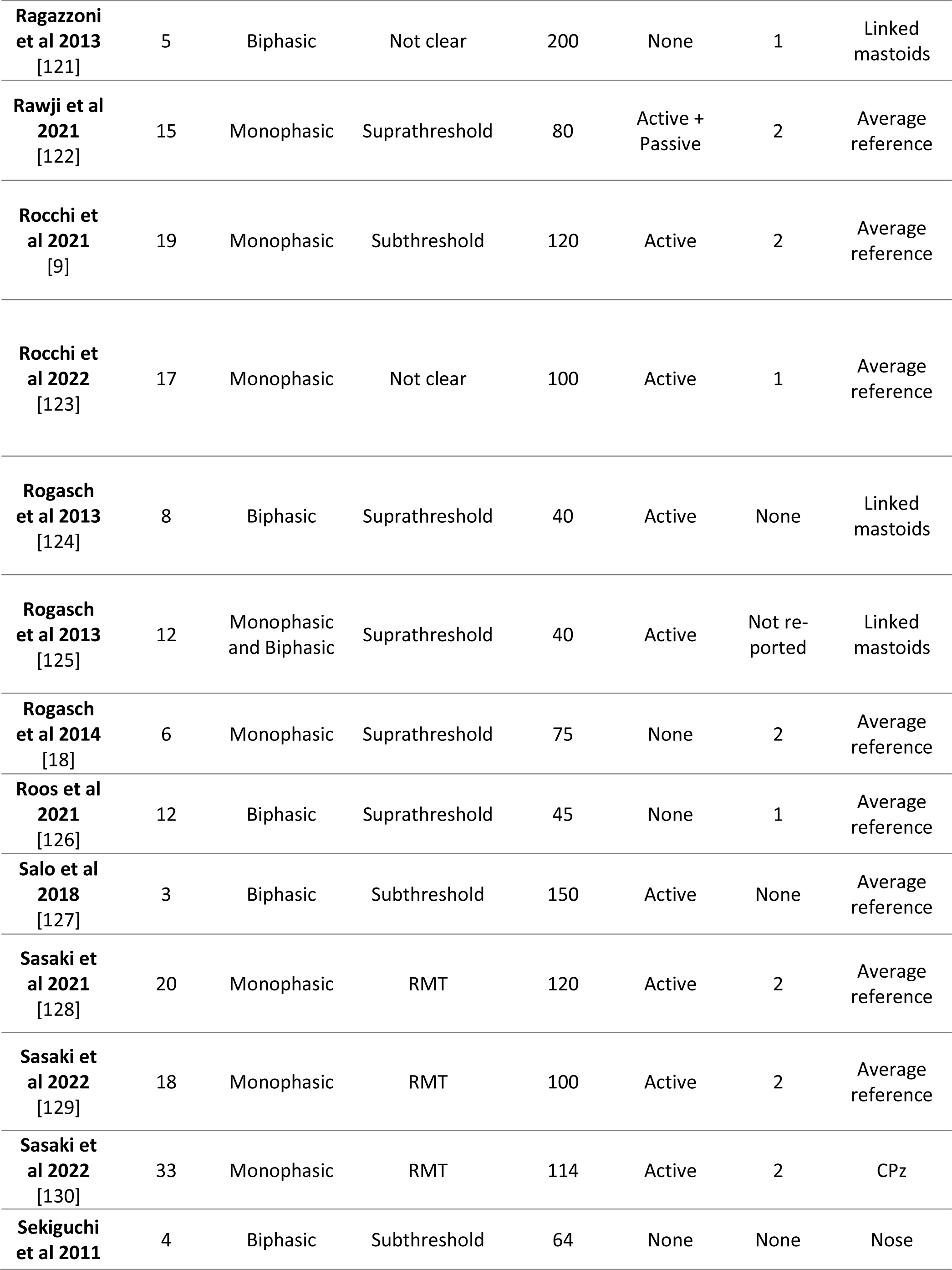
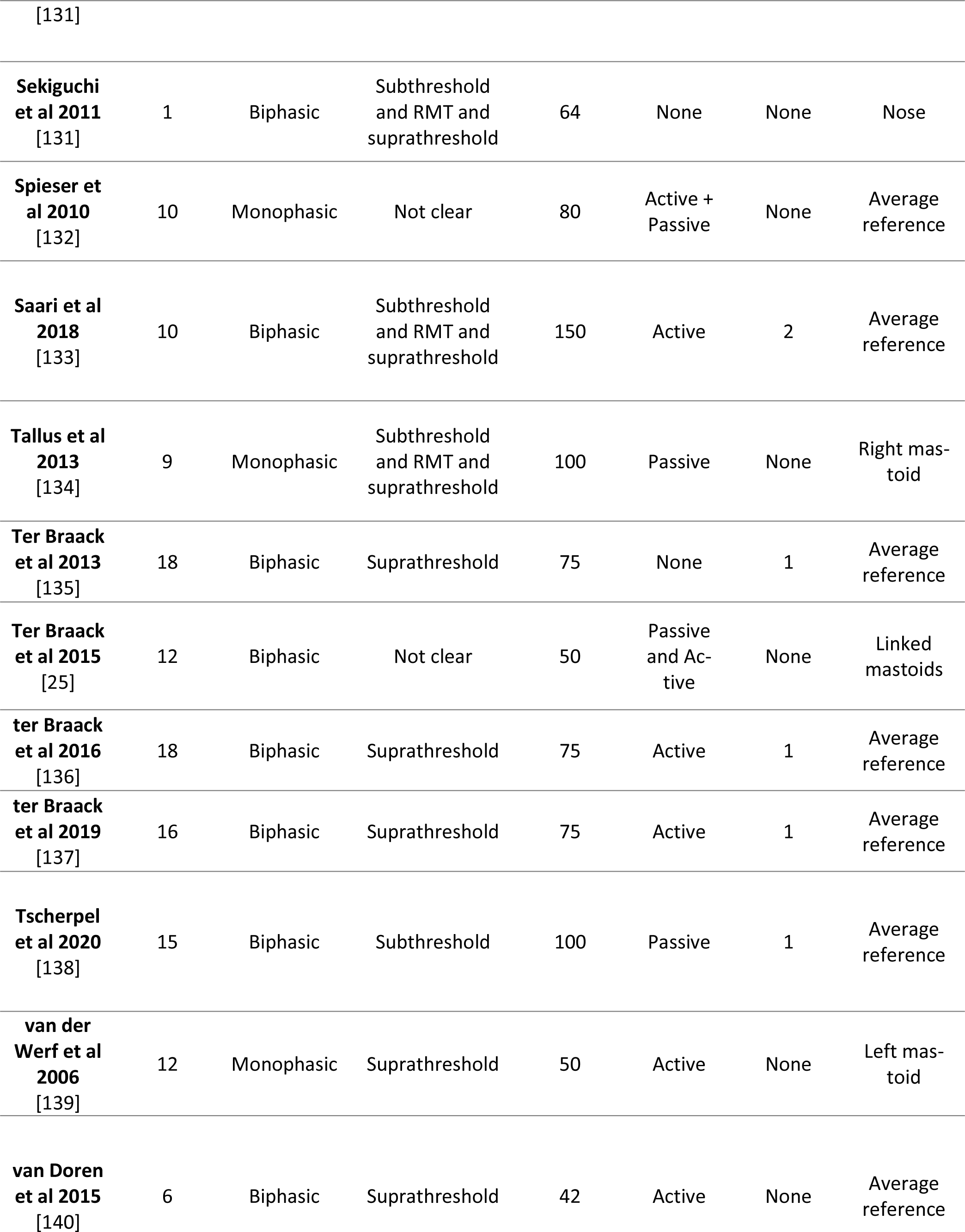
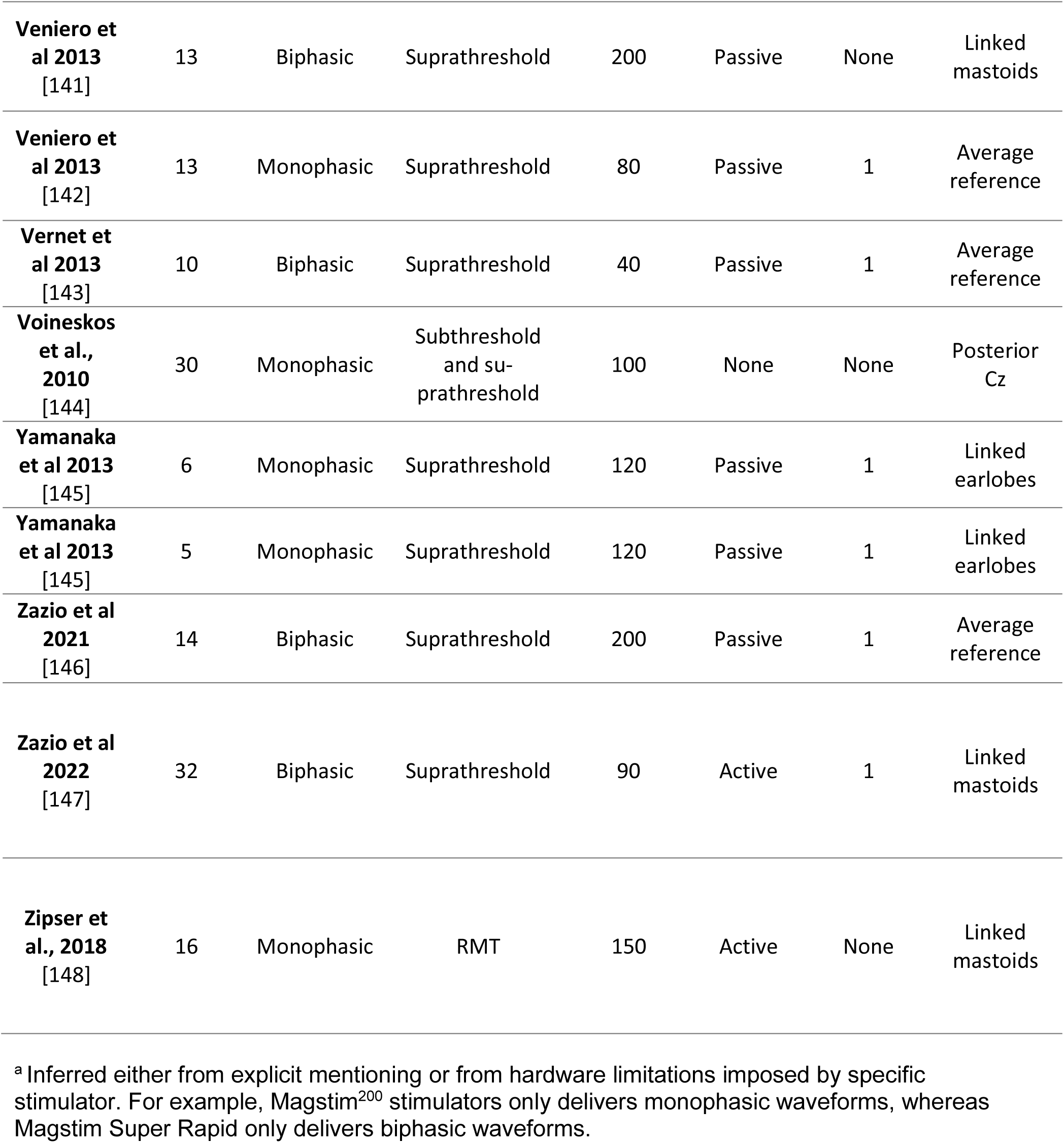
Included studies and a subset of used methods used in these studies.

## Acknowledgements

This work was supported by a “grand solutions” grant “Precision Brain-Circuit Therapy - Precision-BCT)” from Innovation Funds Denmark to Hartwig R. Siebner (grant nr. 9068-00025B) and a collaborative project grant “ADAptive and Precise Targeting of cortex-basal ganglia circuits in Parkinsońs Disease - ADAPT-PD” from Lundbeckfonden to Hartwig R. Siebner (grant nr. R336-2020-1035). Mikkel M. Beck is funded by a post doc grant from the Research fund of the Capital Region Denmark (Region H). Lasse Christiansen holds a postdoc grant from the Lundbeckfonden (R322-2019-2406). Leo Tomasevic holds an ‘Experiment grant’ from Lundbeckfonden (R346-2020-1822).

## Author Contributions (CRediT author statement)

**Conceptualization:** MMB, HRS; **Methodology:** MMB, MH, HRS; **Validation:** MH, LM **Formal analysis:** MMB; **Investigation**: MMB, MH, LM; **Writing-Original draft:** MMB, LC, LT, HRS; **Writing-Review and editing:** MMB, MH, LM, MCV, LC, LT, HRS; **Visualization:** MMB; **Supervision:** LC, LT, HRS; **Funding Acquisition:** MMB, HRS

