## Supplementary material for "Methodological choices matter: A systematic comparison of TMS-EEG studies targeting the primary motor cortex"

### 1.1 Variables extracted from full-text review

We extracted a variety of different variables from the included published studies. In Table S1 below, we present an overview of all the extracted variables per category.

### Table S1. Extracted variables from included studies

| Sample demographics | Stimulation parameters | Experimental setup | EEG acquisition | EEG preprocessing | Data reporting |
|---|---|---|---|---|---|
| Number of subjects | Multiple hemispheres | Type of auditory masking | EEG amplifier manufacturer | High pass filter offline | Time window for reporting |
| Biological sex | Hemisphere stimulated | Type of PEP reduction | EEG amplifier | Low pass filter offline | Electrodes for reporting |
| Age range | Site of stimulation | Type of sham control | EEG amplifier type | Type of filter | Readout reported |
| Age | Determination of stimulation site | Type of physchophysical assessment | Sampling rate | Filter info | Reported high-pass filter |
| | Muscle of interest | Type of task | Number of EEG electrodes | Notch filter | Reported low-pass filter |
| | Method for coil positioning | Type of state control | Placement of electrodes | Software used for preprocessing | Reported reference |
| | Number of stimulations | Type of subject instruction | Position of electrodes | Window for epoching | |
| | Multiple intensities | | EOG | Number of removed epochs | |
| | Determination of intensity | | Type of electrodes | Method used to discard epochs | |
| | Intensity | | Electrode impedance | Baseline correction interval | |
| | Interstimulus interval | | Position of reference electrode | Downsampling | |
| | Jitter | | Position of ground electrode | Re-referencing | |
| | Coil shape | | High pass filter recording (analogue filter) | Window for removing TMS artefact | |
| | Coil diameter | | Low pass filter recording (analogue filter) | Type of interpolation | |
| | Coil name | | Recording software | Specific data analysis tools for TMS-EEG | |
| | Coil orientation | | | ICA | |
| | Stimulator vendor | | | Type of ICA | |
| | Stimulator | | | Number of ICAs performed | |
| | Pulse waveform | | | Number of ICs removed | |
| | Pulse duration | | | Method used to discard ICs | |
| | Pulse waveform inferred | | | | |
| | Current direction | | | | |
| | Current direction inferred | | | | |
| | Recharge delay | | | | |

### 1.2 Methodological choices and trends over time

Below, we present additional information on the used methods and their development over time in published TMS-EEG studies targeting M1. As described in the main text, most studies have used current directions with a posterior-to-anterior directed current (96%). Twenty-three (23%) percentage of studies use a layer of either foam or plastic between the coil and the electrodes to reduce bone conduction of sound and vibration. A few studies (7%) assess participants' percepts of the TMS pulses using either visual analog scales or questionnaires. Most studies performed recordings while subjects are at rest (88%) and a large variety of high-pass and low-pass filtering is being used for preprocessing the data offline after acquisition. As pointed out in the main text, studies mainly reported data from electrodes close to the site of stimulation, but the vertex electrode Cz was also a common choice (Fig S2).

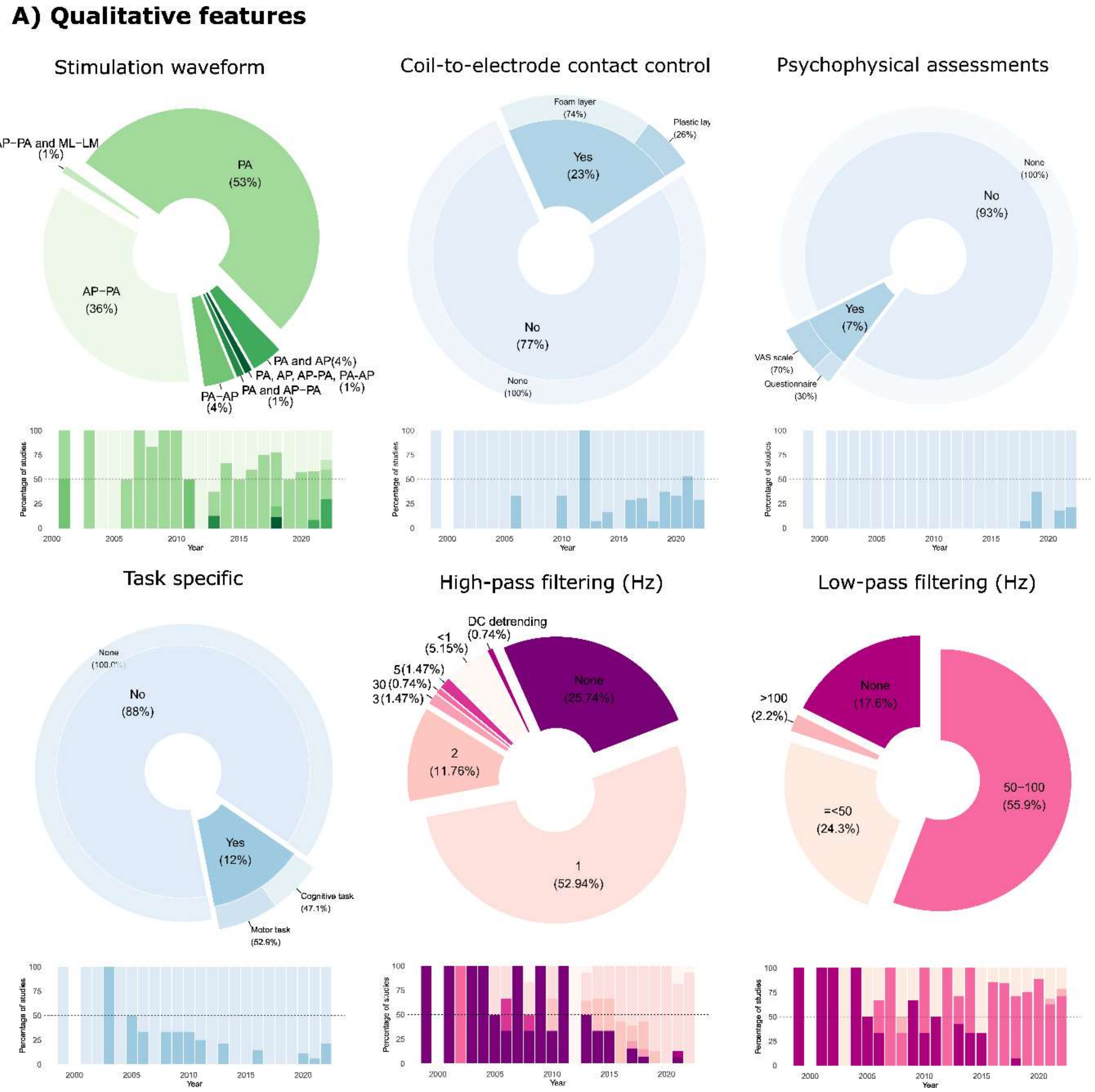


**Figure S1. Methodological choices and development over time.** Displays the relative distribution of used methods in published TMS-EEG studies targeting the M1 and how this has changed over time. Variables relating to TMS parameters (green colors), acquisition parameters (blue colors) and preprocessing strategies (red colors) are displayed.

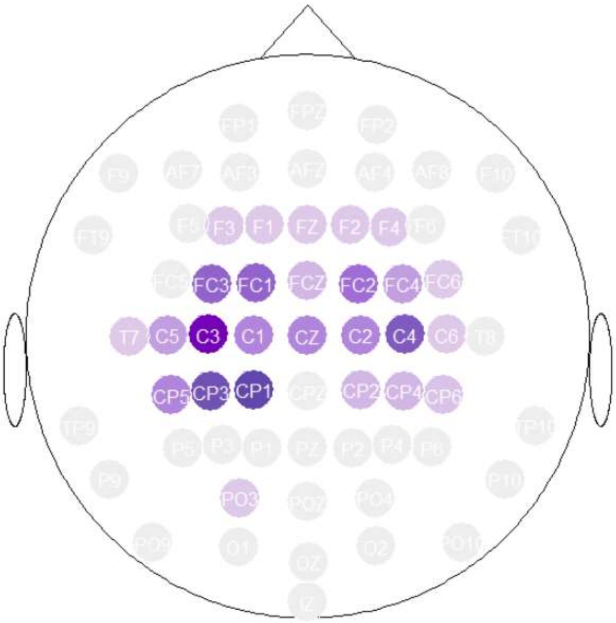


**Figure S2. EEG electrodes from which TEPs were reported from in TMS-EEG studies targeting M1.** The montage shows the most frequently used for reporting TEPs in TMS-EEG experiments regardless of reference electrode and TMS stimulation site. Darker purple colors represent greater proportion of studies reporting from specific electrode. Electrodes around the site of stimulation were most frequently reported. The Cz electrode was also a common choice.

### 1.3 Effects of methodological choices on spontaneous activity

To evaluate whether the differences observed in TEP amplitudes reported in the main text could be explained by general differences in signal amplitudes across used methods, we extracted signal amplitude values from previously published studies at two timepoints prior to the delivery of TMS (-50ms and -10ms with respect to the TMS pulse). We found no significant differences in the averaged, rectified baseline amplitude across studies using different stimulation intensities ($F_{(2,63)}$=0.15, P=0.86), noise masking procedures ($F_{(2,66)}$=0.40, P=0.67), or ICA strategies ($F_{(2,65)}$ =1.35, P=0.27)(Fig S3). This suggests that the differences observed in TEPs were likely not explained by a general difference in EEG signal amplitudes across studies.

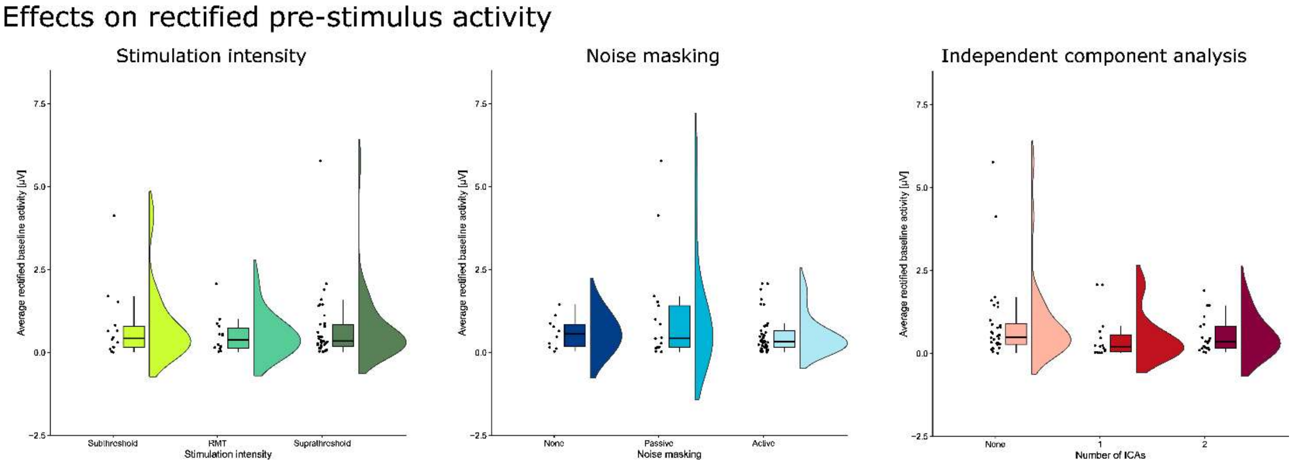


**Figure S3. No differences in pre-stimulus activity across methodological choices.** Displays average rectified amplitudes for spontaneous (baseline) pre-stimulus data across stimulation intensities (left), noise masking (middle) and independent component analysis (right). No statistical differences were observed for any of the conditions.

#### 1.4 Effects of methodological choices on TEP peaks normalized to relative sample size

Previous studies included a different number of subjects. In what follows, we have compared TEP features between studies using different stimulation intensities, noise masking procedures, and independent component analysis strategies after weighting each study's TEP amplitudes by proportion of the total sample. That is, individual studies that included more participants constituting a greater proportion of the total sample size across all studies weighted more compared to studies including fewer participants. For example, a study with 20 subjects received a weight of 20/1269 = 1.58%. This was done as an alternative to inverse variance weighting because measures of variance could not be extracted from most studies. Overall, similar results were found for the TEP features when comparing across different methods after weighting each study:

For stimulation intensity, we found a significant effect on the weighted activation index using a non-parametric Kruskal-Wallis ranked sum test (Chi-squared = 16.35; df = 2; P < 0.001). Suprathreshold stimulation resulted in higher overall activation compared to subthreshold (P = 0.003) and RMT (P = 0.002) stimulation. Although these effects seemed to generalize across multiple TEP peaks, they were largest – or most consistent - for the N45, P60 and N100 peak. No effects were observed on the N100-P180 complex (Chi-squared = 3.42; df = 2; P = 0.18)(Figure S4AB).

For noise masking, we found a significant effect on the weighted 'activation index' (Chi-squared = 8.16; df = 2; P = 0.017). Using active noise masking and refraining from using noise masking resulted in lower overall activation than using passive noise masking (P = 0.04 and P = 0.008, respectively). A significant effect was also observed for the N100-P180 complex (Chi-squared = 11.90; df = 2; P = 0.003), resulting from a lower weighted amplitude observed between studies using passive noise masking and studies using no noise masking (P = 0.007) or active noise masking (P = 0.009) (Figure S4AB).

For independent component analysis, lower values were seen for the weighted 'activation index' after using ICA compared to not using ICA. However, there was no significant effect on the 'activation index' (Chi-squared = 2.91; df = 2; P = 0.23). In line with the results presented in the main text, we did find a significantly smaller N100-P180 complex following two rounds of ICA compared to no ICA (P = 0.02) (Figure S4AB).

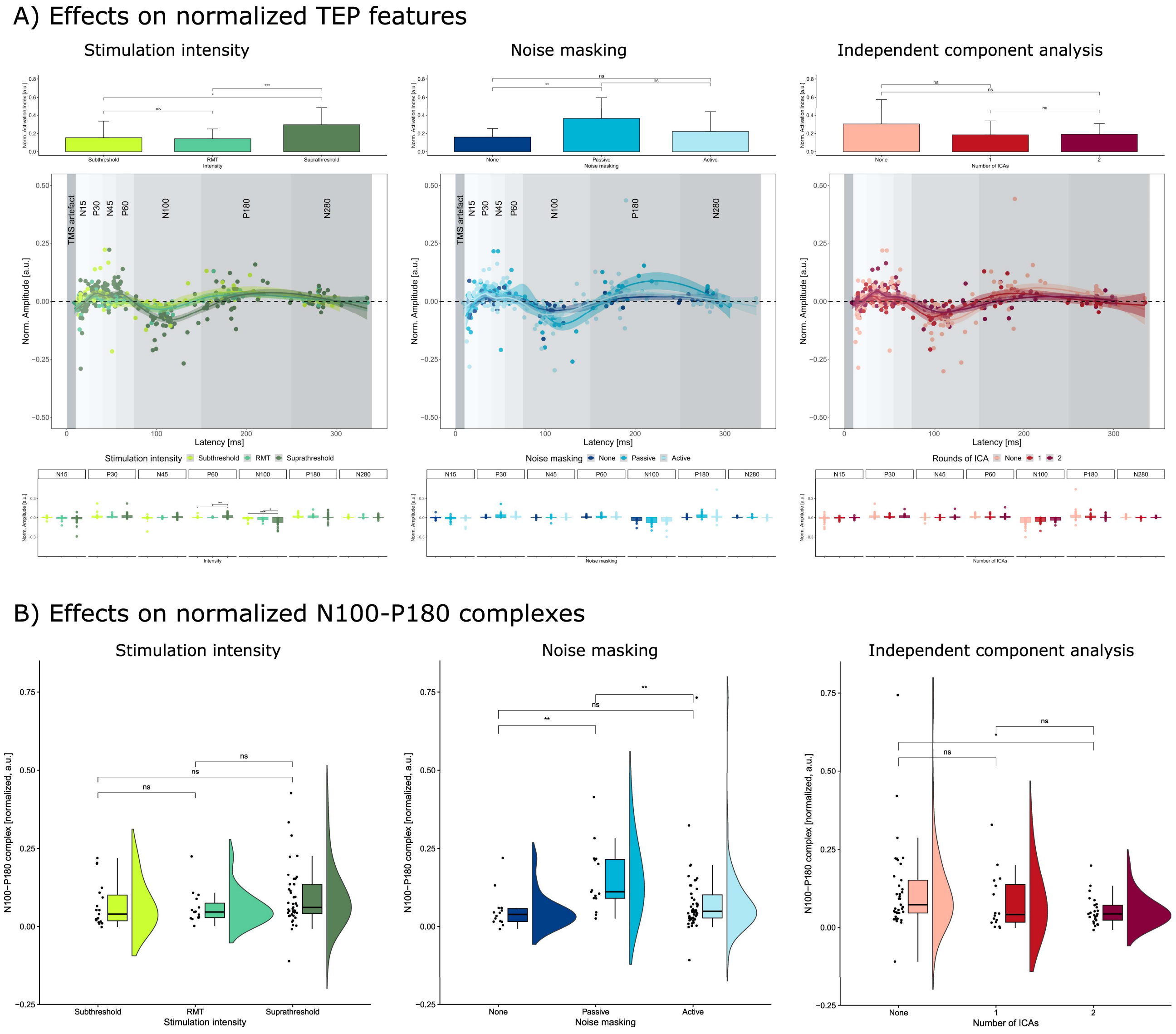


**Figure S4. The impact of methodological choices on sample size normalized transcranial evoked EEG potentials.** (A) Effects on total activation and TEP peaks. For each subfigure, the overall 'activation index', computed as the sum of the absolute peak values for each TEP peak, is presented in the top row. The middle row displays TEP waveforms reconstructed from extracted latencies (x-axis) and amplitudes (y-axis) and the bottom row displays a bar-graph of average values of different prototypical peaks for each of the methodological conditions. Individual values have been normalized to the relative sample size in the study from which it was extracted. The leftmost figure displays the impact of stimulation intensity all other parameters aside. Suprathreshold stimulation leads to a greater overall 'activation index' across studies. The P60 and N100 display greater amplitudes for suprathreshold stimulation compared to subthreshold and threshold intensities. The middle figure displays the impact of noise masking all other parameters aside. Studies using passive noise masking result in a higher overall 'activation index' and this effect seems to be general across various peaks. The rightmost figure displays the impact of using independent component analysis for preprocessing all other parameters aside. Studies using ICA generally result in a smaller overall 'activation index', but this difference was not statistically significant. (B) Effects on N100-P180 complex. The leftmost figure displays the impact of stimulation intensity on normalized TEPs. No statistical effects were observed. The middle figure displays effect of noise masking procedures on N100-P180 complexes. Using passive noise masking resulted in higher N100-P180 complexes compared to active and no noise masking. The rightmost figure displays the impact of using ICA for preprocessing. Using two rounds of ICA resulted in lower N100-P180 complexes compared to not using ICA.

1.5 **Dependence between use of methods for main effect analyses**

In the main body of this work, we compared TEP peak amplitudes across already published studies using different methods. These findings are equivalent to main effects and should be interpreted as such. However, it is possible that certain methodological choices depend on one another, i.e., they might be nested in one another. In Figure S5, we explored this possibility for the methodological choices investigated here. We did not find clear dependencies between the use of different methods.

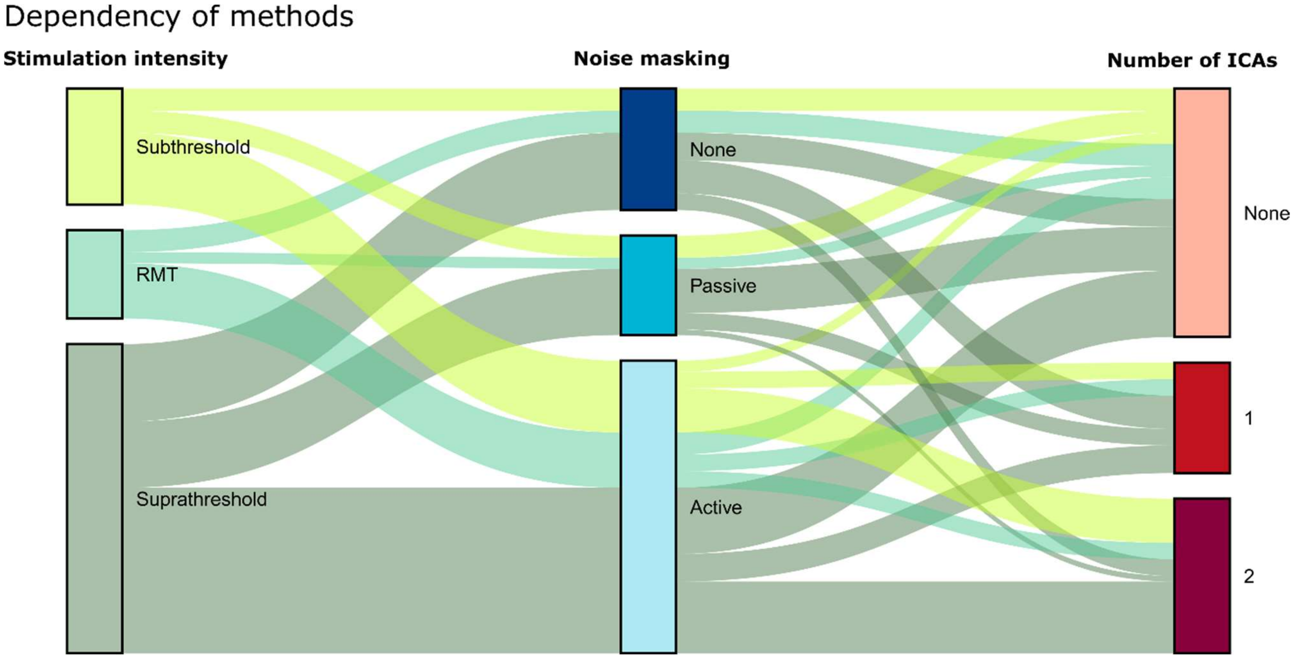


**Figure S5. A parallel-set plot of dependency between methodological choices.** Displays the proportion of studies using different combinations of methods. No clear dependencies were observed for the parameters that we have explored in the main text of this work.
